# Dysregulated expression of inflammasome and extracellular matrix genes in *C9orf72*-ALS/FTD microglia

**DOI:** 10.1101/2025.02.13.638162

**Authors:** Nisha S Pulimood, Louise Thiry, Ye Man Tang, Stefano Stifani

**Affiliations:** Department of Neurology and Neurosurgery, Montreal Neurological Institute-Hospital, McGill University, 3801, rue University, Montreal, QC H3A 2B4 Canada

**Keywords:** amyotrophic lateral sclerosis, C9orf72, extracellular matrix, inflammasome, microglia, RNA sequencing

## Abstract

Hexanucleotide repeat expansion (HRE) in the non-coding region of the gene *C9orf72* is the most prevalent mutation in amyotrophic lateral sclerosis (ALS) and frontotemporal dementia (FTD). The *C9orf72* HRE contributes to neuron degeneration in ALS/FTD through both cell-autonomous mechanisms and non-cell autonomous disease processes involving glial cells such as microglia. The molecular mechanisms underlying the contribution of *C9orf72*-HRE microglia to neuron death in ALS/FTD remain to be fully elucidated. In this study, we generated microglia from human *C9orf72*-HRE and isogenic iPSCs using three different microglia derivation methods. RNA sequencing analysis reveals a cell-autonomous dysregulation of extracellular matrix (ECM) genes and genes involved in pathways underlying inflammasome activation in *C9orf72*-HRE microglia. In agreement with elevated expression of inflammasome components, conditioned media from *C9orf72*-HRE microglia enhance the death of *C9orf72*- HRE motor neurons implicating microglia-secreted molecules in non-cell autonomous mechanisms of *C9orf72* HRE pathology. These findings suggest that aberrant activation of inflammasome-mediated mechanisms in *C9orf72*-HRE microglia results in a pro-inflammatory phenotype that contributes to non-cell autonomous mechanisms of motor neuron degeneration in ALS/FTD.

**Summar**y**:** This study describes phenotypic alterations in *C9orf72*-ALS/FTD microglia implicating extracellular matrix remodeling and inflammasome activation in microglia-mediated neurodegeneration in ALS/FTD.

## Introduction

Amyotrophic lateral sclerosis (ALS) is an incurable motor neuron disease characterized by neuronal death in the cerebral cortex, brain stem, and spinal cord. ALS usually starts during adulthood and causes progressive paralysis leading to the death of most sufferers within 2 to 5 years following diagnosis, mainly due to respiratory failure. ALS also affects several non-motor systems and subcortical structures, and cognitive and behavioral impairments similar to frontotemporal dementia (FTD) symptoms are present in up to 50% of ALS patients (Benatar et al., 2024; Kirola et al., 2022).

The majority of ALS cases are sporadic, implying that the mechanisms triggering motor neuron loss are not exclusively inherited. However, approximately 10%-15% of ALS cases are transmitted in families: these cases are referred to as familial ALS. Multiple deleterious variants in numerous genes either drive motor neuron degeneration, increase susceptibility to the disease, or influence the rate of disease progression. Several ALS-mutated genes have a negative impact on shared cellular processes, including, but not limited to, RNA metabolism and protein homeostasis, nucleocytoplasmic trafficking, endoplasmic reticulum stress, dynamics of ribonucleoprotein bodies, mitochondrial functions, and autophagy (Goutman et al., 2022; Nijs and Van Damme, 2024).

Intronic hexanucleotide (GGGGCC – ‘G_4_C_2_’) repeat expansions (HRE) in the gene *chromosome 9 open reading frame 72* (*C9orf72*) are the most frequent genetic cause of familial ALS/FTD (DeJesus-Hernandez et al., 2011; Renton et al., 2011). In contrast to healthy individuals, who have, on average, 2-30 hexanucleotide repeats, ALS/FTD patients have *C9orf72* expansions with hundreds and even thousands of repeats. There are three (not mutually exclusive) proposed mechanisms to explain the effects of the *C9orf72* HRE: partial loss (haploinsufficiency) of *C9orf72* function caused by decreased expression of the C9orf72 mRNA and protein; toxic gain of function from the accumulation of sense G_4_C_2_ and antisense G_2_C_4_ repeat-containing RNA from bidirectionally transcribed expansions; and toxic gain of function from dipeptide repeat proteins aberrantly generated by repeat-associated non-ATG translation (Balendra and Isaacs, 2018; Gendron et al., 2018). The combination of loss- and gain-of-function mechanisms associated with the *C9orf72* HRE have wide-ranging consequences in multiple cell types, in agreement with the broad expression pattern of the *C9orf72* gene and the localization of the C9orf72 protein to multiple cellular compartments. Multiple cellular processes are affected in *C9orf72*-mutated motor neurons leading to altered mitochondria, oxidative stress, impaired autophagy, and impaired RNA metabolism, among other phenotypes (Chew et al., 2015; de Calbiac et al., 2024; Donnelly et al., 2013; Lopez-Gonzalez et al., 2016; Metha et al., 2021; Tran et al., 2015).

Like other neurodegenerative diseases, ALS is characterized by extensive neuroinflammation involving astrogliosis, activation of microglia, and infiltration of peripheral immune cells at sites of neuronal degeneration. Glial cells have a significant impact on disease onset and progression by mediating non-cell autonomous disease mechanisms (Appel et al., 2021; Calma et al., 2024; Izrael et al., 2020; Majewski et al., 2025). Intercellular communication between motor neurons and microglia, the resident immune cells of the central nervous system, play an important role in the degeneration of motor circuits in ALS. Microglia have the potential to perform both neuroprotective and neurotoxic roles in ALS, depending on their activation states in response to the local microenvironment. During initial phases of the disease microglia can protect motor neurons by performing scavenging, anti-inflammatory, and neuroprotective functions. During disease progression, however, with rising oxidative stress and injury within motor neurons, microglia can contribute to motor-neuron death by switching to pro-inflammatory and neurotoxic phenotypes (Beers and Appel, 2019; Majewski et al., 2025; Mimic et al., 2023). Much remains to be learned about the mechanisms underlying the dynamic intercellular communication between motor neurons and microglia in ALS.

*C9orf72* is robustly expressed in microglia and other myeloid-lineage cells. Analysis of post-mortem brains from *C9orf72*-ALS/FTD patients revealed morphological and immunological signs suggestive of microglia activation (Lant et al., 2014), as did *ex vivo* diffusion tensor MRI studies of tissues from ALS patients including *C9orf72* cases (Cardenas et al., 2017). Evidence that *C9orf72* is required for the normal function of myeloid cells first came from studies in mice lacking the murine *C9orf72* ortholog in all tissues. *C9orf72*^-/-^ (homozygous knockout) mice develop and age normally but exhibit several immune cell phenotypes, including lysosomal accumulation and altered immune responses in macrophages and microglia, with associated age-related neuroinflammation (Atanasio et al., 2016; Burberry et al., 2016; O’Rourke et al., 2016; Sudria-Lopez et al., 2016). Studies in which the human *C9orf72* HRE was introduced into mice showed signs of pathological expression of *C9orf72* HRE in the cortex including the appearance of hallmarks of microglia activation (Nicholson et al., 2018; Zhang et al., 2018). Transcriptomic analysis of *C9orf72* knockout mouse microglia revealed gene expression changes consistent with an inflammatory phenotype and correlated with increased microglia-mediated synaptic loss, which together imply that loss of *C9orf72* results in a proinflammatory and neurotoxic microglia phenotype (Lall et al., 2021). More recent studies using microglia generated from induced pluripotent stem cells (iPSCs) derived from ALS/FTD patients with *C9orf72* amplifications have shown that human *C9orf2*-HRE microglia exhibit typical hallmarks of *C9orf72* HRE pathology such as G_4_C_2_-repeat RNA foci, expression of dipeptide repeat proteins, and decreased expression level of C9orf72 (Lazzarini et al., 2023; Vahsen et al., 2023). Moreover, lipopolysaccharide (LPS)-primed *C9orf72*-HRE microglia were observed to have a pro-inflammatory phenotype and to be toxic to motor neurons when compared to healthy and/or isogenic microglia as controls (Banerjee et al., 2023; Vahsen et al., 2023).

Recent studies in both animal and human cell models have shown that the *C9orf72* HRE is associated with activation of the Nucleotide-Binding-Oligomerization Domain (NOD)-and Leucine-Rich Repeat (LRR)-containing (NLR) family, Pyrin-Domain-Containing 3 (NLRP3) inflammasome in microglia (Banerjee et al., 2023; Fu et al., 2023; Rivers-Auty et al., 2024; Shu et al., 2023; Trageser et al., 2023). The NLRP3 inflammasome is a cellular immune sensor that mediates pathological inflammation in autoimmune, metabolic, malignant, and neurodegenerative conditions. It detects stress signals, such as tissue damage or pathogen invasion, via immune cell components referred to as pattern recognition receptors (PRRs). For instance, the PRR Toll-like Receptor 4 (TLR4) can detect ‘danger signals’ and respond by enlisting Nuclear Factor-kappaB (NF-κB) to activate the transcription of genes encoding proteins involved in the NLRP3 inflammasome, including *NLRP3* itself. The NLRP3 protein assembles with other components, such as Apoptosis-associated Speck-like protein containing a Caspase recruitment domain (ASC), to form the active inflammasome complex. The NLRP3 complex activates Caspase-1 (CASP1) which in turn induces the release of inflammatory cytokines like Interleukin-1 beta (IL1B) and IL18 and the activation of pyroptosis, a highly inflammatory form of lytic cell death (Fu and Wu, 2023; Sharma and Kanneganti, 2021). Little is known, however, about the molecular mechanisms underlying NLRP3 inflammasome activation in *C9orf72*-mutated microglia.

In this study, we characterized the transcriptomic changes associated with *C9orf72* amplification in microglia derived from *C9orf2*-HRE iPSCs, and isogenic control iPSCs, using three different microglia differentiation methods. We specifically compared derivation protocols that generate microglia with distinct intrinsic inflammatory profiles to alleviate the possibility that differentiation protocol-specific basal activation of the induced microglia might confound the study of the impact of the *C9orf72* HRE on inflammatory mechanisms. Here we show a consistent dysregulation of several genes involved in pathways underlying NLRP3 inflammasome activation in *C9orf72*-HRE microglia. Numerous dysregulated genes in *C9orf72*-HRE microglia encode extracellular matrix (ECM) proteins, suggesting a link between ECM remodelling and inflammasome activation in microglia in ALS/FTD. We show further that conditioned media from *C9orf72*-HRE microglia enhance the death of *C9orf72*-HRE motor neurons *in vitro*. We hypothesize that cell-autonomous alterations in gene expression associated with inflammasome activation in *C9orf72*-HRE microglia contribute to neuronal death in ALS/FTD. The transcriptomic data generated by this study would serve as an objective source of hypothesis-generating data that may reveal novel mechanisms of microglial involvement in *C9orf72*-mutated ALS/FTD.

## Results

### Generation of C9orf72-mutated human iPSC-derived microglia using three different differentiation protocols

Activation of the NLRP3 inflammasome in microglia was observed in both mouse and human cells harbouring the *C9orf72* HRE (Banerjee et al., 2023; Fu et al., 2023; Rivers-Auty et al., 2024; Shu et al., 2023; Trageser et al., 2023). Little information is available, however, about the identity of genes that are dysregulated as part of the NLRP3 inflammasome activation in *C9orf72*-HRE (*C9orf72*-mutated) microglia. To address this gap in knowledge, we implemented a comprehensive RNA sequencing (RNAseq) study of human microglia derived from *C9orf72*-mutated iPSC lines, and matching isogenic lines. Four previously studied (Chi et al., 2023; Ho et al., 2021; Gao et al., 2024; Thiry et al., 2024) iPSC lines were used for microglia derivation: the *C9orf72*-mutated iPSC lines CS52iALS-C9n6 (CS52-C9n6-M hereafter) and CS29iALS-C9n1 (CS29-C9n1-M), and their matching isogenic (ISO) control lines CS52iALS-C9n6.ISOxx (CS52-C9n6-ISO) and CS29iALS-C9n1.ISOxx (CS29-C9n1-ISO). Previous studies have demonstrated the presence of *C9orf72* HRE RNA foci and dipeptide repeat expression in neuromuscular organoids generated using these *C9orf72*- mutated iPSC lines (Gao et al., 2024), providing evidence that they can recapitulate *C9orf72* HRE-associated pathological features *in vitro*.

We generated iPSC-derived microglia using three separate differentiation methods based on previously published protocols (Douvaras et al., 2017; Haenseler et al., 2017; McQuade et al., 2018). The use of multiple derivation protocols was aimed at mitigating the confounding effects that might result from the variability in the basal properties of iPSC-derived microglia generated using different *in vitro* differentiation methods (Tang et al., 2022). Hereafter, we shall operationally refer to the three microglia differentiation protocols used in the present study as protocols P1 (Douvaras et al., 2017), P2 (McQuade et al., 2018), and P3 (Haenseler et al., 2017) (Figure 1A–C). With all protocols, we allowed microglia to acquire a mature phenotype *in vitro* by differentiating the cells for at least as long as indicated in the original publications and then confirming robustness and maturity of the induced cultures as we have previously described (Tang et al., 2022).

**Figure 1.**
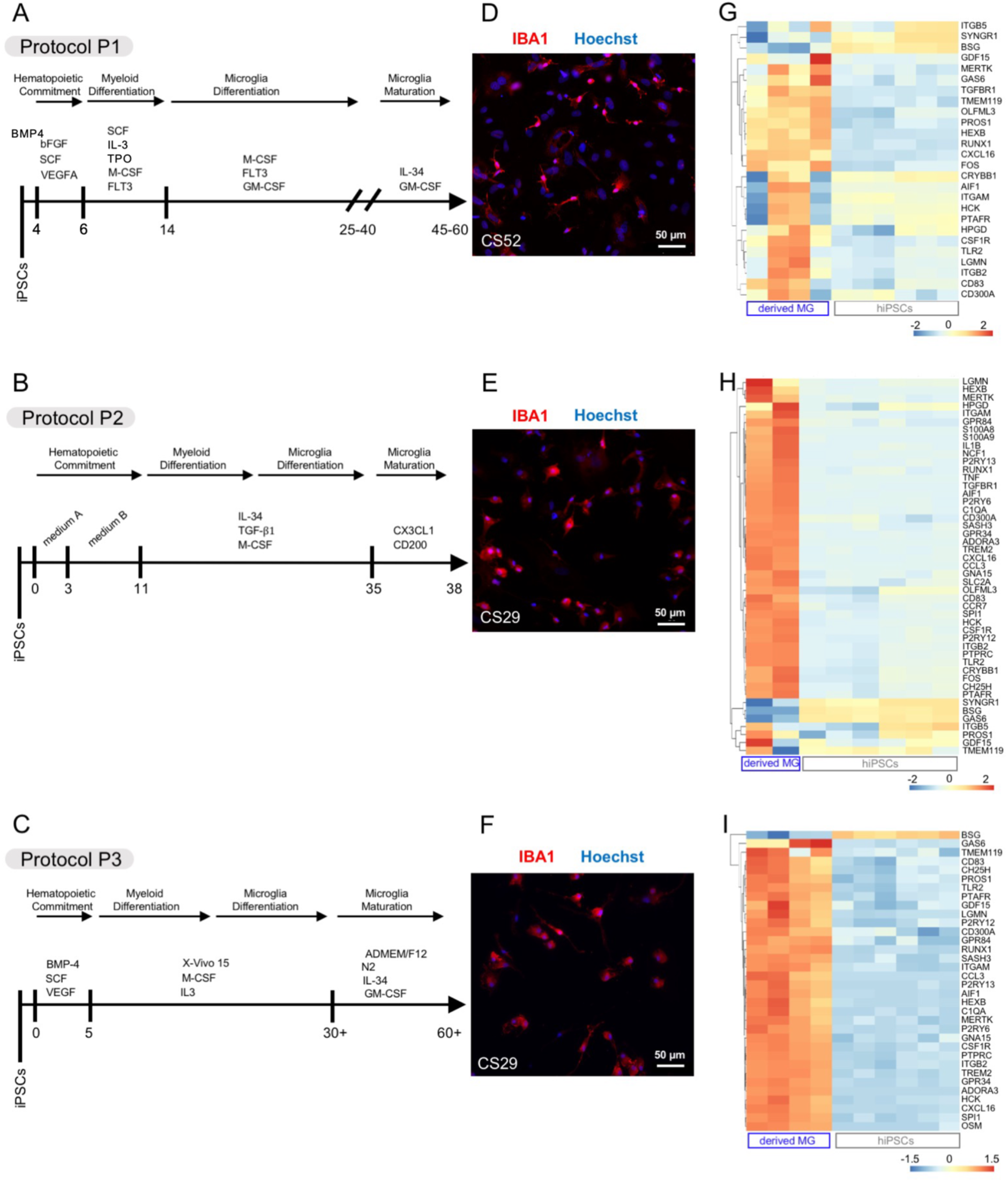
Generation and characterization of iPSC-derived microglia using three separate methods. (A–C) Schematics of three microglia differentiation protocols, referred to as “protocol P1” adapted from Douvaras et al. (A), “protocol P2” adapted from McQuade et al. (B), and “protocol P3” adapted from Haenseler et al. (C). (D–F) Representative images of microglia preparations differentiated from iPSC lines CS52-C9n6-ISO (CS52) and CS29-C9n1-ISO (CS29) with protocol P1 (D), P2 (E), or P3 (F) and immunostained with anti-IBA1 antibody (red) and counterstained with Hoechst (blue). Scale bar, 50 μm. (G–I) Heatmaps showing known microglia/macrophage marker genes expression in undifferentiated iPSCs or microglia preparations differentiated from the CS52 and CS29 iPSC lines with protocol P1 (G), P2 (H) or P3 (I). Each column corresponds to one sample (undifferentiated iPSCs (hiPSCs) or isogenic iPSC-derived microglia (derived MG), as indicated at the bottom, and each line corresponds to one gene. Subgroups of genes with correlated expression patterns are portrayed by the dendrogram on the left side of each heatmap. Relative levels of gene expression are indicated by a color scale, with cool colors corresponding to low expression and warm colors corresponding to increased/high expression. Hierarchical clustering of rows was applied to the heatmaps to group genes with correlated expression patterns, resulting in different gene lists, depending on the samples analyzed in the given heatmap. The gene list used for this analysis was as follows: *HEXB, P2RY12, CXC3CR1, P2RY13, TREM2, S100A8, TMEM119, S100A9, RNASE4, GPR34, FCRLS, SIGLECH, OLFML3, FOS, SCLO2B1, TGFBR1, SLC2A5, CAMP, ITGB5, CRYBB1, SYNGR1, GPR56, NGP, CMRF35, HPGD, P2RY12, GPR34, MERTK, C1QA, PROS1, GAS6, ITGAM, ITGB2, CSF1R, PTPRC, AIF1, ADORA3, LGMN, GPR84, CCR7, BCL2A1D, TNF, NCF1, GDF15, OSM, LMC25, H2-OA, CD83, CCL3, GNA15, IL1B, PFAU, CCL9, TMEM119, C1QA, LRF8, CXCL16, CH25H, HCK, CCL12, PTAFR, CD300A, LRT5, STPI1, SELPIG, SASH3, BSG, TLR2, P2RY6, CDI4, BCL2A1A, BCL2A1C, RUNX1, SPI1*. Abbreviations: hiPSC, human iPSC; MG, microglia.

Examining isogenic iPSC lines CS52-C9n6-ISO and CS29-C9n1-ISO, we observed that all three microglia derivation protocols generated cultures highly enriched in cells expressing typical microglia/macrophage markers, including the protein IBA1, by immunocytochemistry (Figure 1D–F). More importantly, RNAseq data showed that the majority of 49 previously characterized microglia/macrophage cell marker genes (Butovsky et al., 2014; Hickman et al., 2013; Muffat et al., 2016; Zhang et al., 2014) were upregulated in microglia generated with all three protocols, when compared to separate undifferentiated iPSC lines (Figure 1G–I).

Microglia generated using protocol P2 had the highest number of significantly upregulated genes encoding pro-inflammatory molecules, such as Chemokine (C-C motif) ligand 3 (CCL3), C-C Motif Chemokine Receptor 7 (CCR7), C-X-C Motif Chemokine Ligand 16 (CXCL16), G Protein Coupled Receptor 84 (GPR84), Hematopoietic Cell Kinase (HCK), IL1B, S100 Calcium Binding Protein A9 (S100A9), S100 Calcium Binding Protein B (S100B), Tumor Necrosis Factor (TNF), when compared to microglia generated with protocols P1 and P3 (Figure 1I-G). Consistently, direct comparison of the expression of the same 49 microglia/macrophage genes across a panel of RNAseq datasets from microglia generated using the three different methods showed that protocol 2-microglia had the highest levels of pro-inflammatory genes. Protocol P3 had the lowest levels of these genes, suggesting that method 3-microglia are the least developmentally mature and/or least activated compared to microglia obtained using the other two protocols, even after longer *in vitro* differentiation times (Figure 2). Together, these observations suggest that microglia generated using method P2 are more pro-inflammatory than microglia obtained using protocols P1 and P3 under comparable experimental conditions.

**Figure 2.**
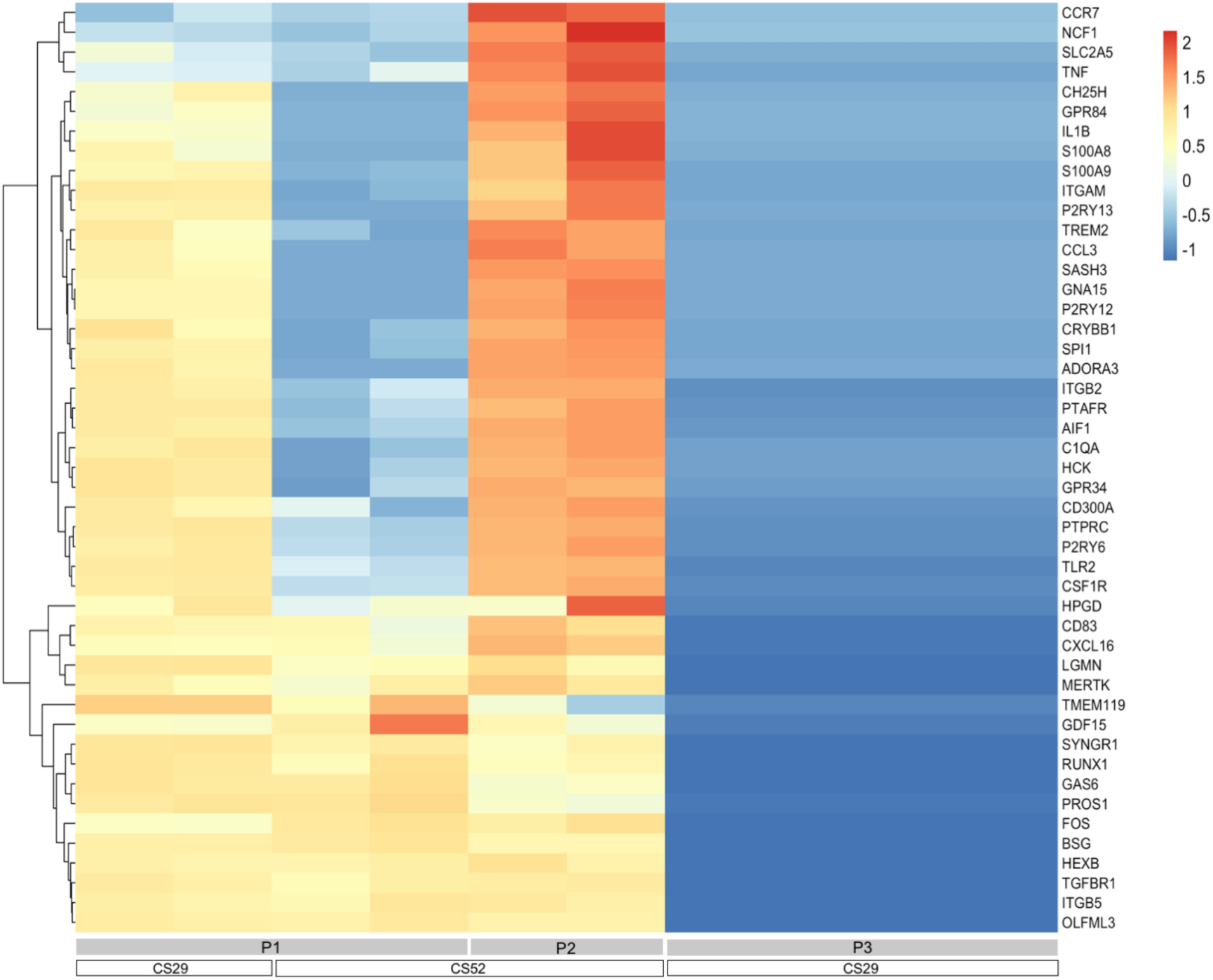
Comparison of microglial/macrophage genes expression in iPSC-derived microglia obtained with three different protocols. Heatmap showing known microglia/macrophage marker genes expression in microglia preparations differentiated from the CS52-C9n6-ISO (CS52) and CS29-C9n1-ISO (CS29) iPSC lines with protocols P1, P2, or P3. Each column corresponds to one isogenic sample (generated from the CS52 or the CS29 iPSC line with protocols P1, P2 or P3, as indicated at the bottom) and each line corresponds to one gene. Subgroups of genes with correlated expression patterns are portrayed by the dendrogram on the left side of the heatmap. Relative levels of gene expression are indicated by a color scale, with cool colors corresponding to low expression and warm colors corresponding to increased/high expression. The gene list used for this analysis was the same as the list shown in the legend to Figure 1.

### Dysregulated transcriptome of C9orf72-mutated human iPSC-derived microglia

All iPSC lines, *C9orf72*-mutated and isogenic, were used to generate microglia using at least two of the three derivation methods described above. All microglia preparations (25 in total) were analysed by RNAseq. To search for cell-intrinsic transcriptomic changes associated with the *C9orf72* HRE, all microglia preparations were not exposed to lipopolysaccharide (LPS) or other treatments inducing microglia activation prior to RNA collection.

We first analyzed RNA isolated from three separate microglia cultures derived from the *C9orf72*-mutated iPSC line CS52-C9n6-M and three microglia cultures from the matching isogenic control line CS52-C9n6-ISO. These 6 biological samples were obtained using differentiation protocol P1, which in our hands generates microglia that are less intrinsically pro-inflammatory than microglia generated using method 2 (Figure 2). Principle component analysis showed that CS52-C9n6-M microglia grouped together along one dimension, clustering away from CS52-C9n6-ISO microglia (Figure 3A). Comparing the transcriptomic data from these six separate preparations using an EdgeR (version 4.0) analysis pipeline for differentially expressed gene (DEG) analysis (Chen *et al*., 2016), we identified 614 genes that were significantly upregulated in *C9orf72*-mutated microglia, while 165 genes were downregulated (adjusted p-value (false discovery rate; FDR) <0.05, with no fold-change cutoff) (Figure 3B). The top 30 most upregulated DEGs in CS52-C9n6-M microglia included several genes that have previously been associated with ALS, such as *Cluster of Differentiation 14* (*CD14*), *Class II Major Histocompatibility Complex Transactivator* (*CIITA*), *Fc Gamma Receptor IIb* (*FCGR2B*), *Major Histocompatibility Complex, Class II, DQ Alpha 1* (*HLA- DQA1*), and *Colony Stimulating Factor 1 Receptor* (*CSF1R*) (Beers et al., 2022; Chi et al., 2023; Martinez-Muriana et al., 2016; Sanfilippo et al., 2017; Xie et al., 2021) (Figure 3C). Several genes involved in microglia/macrophage activation were also present within the top 30 DEGs, including *Cluster of Differentiation 74* (*CD74*), *Macrophage Scavenger Receptor 1* (*MSR1*), *Phosphoinositide-3-Kinase Regulatory Subunit 5* (*PIK3R5*), *Phosphatidylinositol-4,5- Bisphosphate 3-Kinase Catalytic Subunit Gamma* (*PIK3CG*), to name a few (Gu et al., 2024; Ji et al., 2022; Lajqi et al., 2019; Peferoen et al., 2015). Consistently, combined Gene Ontology (GO) and Kyoto Encyclopedia of Genes and Genomes (KEGG) analysis of the upregulated DEGs in CS52-C9n6-M microglia generated using protocol P1 revealed enrichment in pathways associated with ‘immune response’, ‘inflammatory response’, ‘innate immune response’, to name a few (Figure 3D). The latter finding resembles the observation by Vahsen and colleagues that LPS-primed *C9orf72*-mutated iPSC-derived microglia (obtained using a protocol resembling protocol P3 in our study) exhibit enrichment in DEGs involved in pathways associated with immune-cell activation (Vahsen et al., 2023).

**Figure 3.**
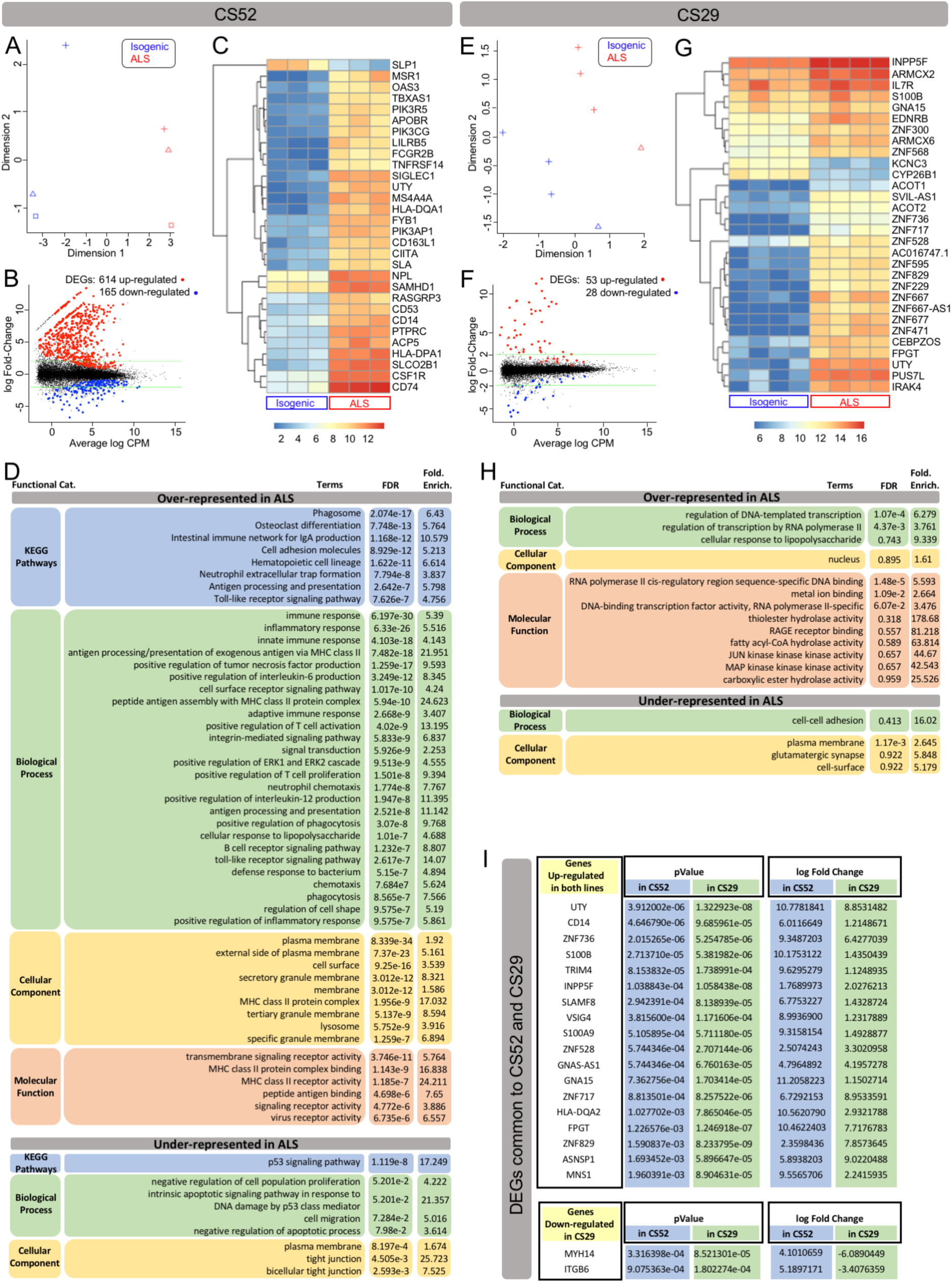
Differential gene expression between corrected isogenic and *C9orf72*-mutated microglia derived using protocol P1. (A–H) Differential gene expression and gene ontology analysis of CS52-C9n6-M versus CS52-C9n6-ISO microglia (CS52 collectively) (A–D) or CS29-C9n1-M versus CS29- C9n1-ISO microglia (CS29 collectively) (E–H). (A and E) Principal component analysis plots showing the relationship between the transcriptomic profiles of CS52 (A) or CS29 (E) iPSC-derived microglia. Each plotted data point represents one biological replicate, and the different symbols refer to the different batches. Blue data points correspond to corrected isogenic samples (Isogenic); red data points correspond to *C9orf72*-mutated samples (ALS). (B and F) Mean-difference plots (MA plots) showing the average log of Count per Million (CPM; X-axis) and log Fold Changes (Y-axis), highlighting the differentially expressed genes (DEGs) in *C9orf72*-mutated (ALS) compared to corrected isogenic microglia differentiated from the CS52 (B) or CS29 (F) iPSC lines. Each plotted data point represents one gene: black data points correspond to genes that are not DEGs, red data points correspond to up- regulated DEGs, and blue data points correspond to down-regulated DEGs (p-adjusted <0.05, with no Fold Change cutoff). The green lines indicate 2-fold up or down. The numbers of up- and down- regulated DEGs are indicated on top of each MA plot. (C and G) Heatmaps of the topmost DEGs in *C9orf72*-mutated (ALS) compared to corrected isogenic microglia differentiated from the CS52 (C) or CS29 (G) iPSC lines. For each heatmap, each column corresponds to one sample (Isogenic or ALS, as indicated at the bottom) and each line corresponds to one gene. Relative levels of gene expression are indicated by a color scale, with cool colors corresponding to low expression and warm colors corresponding to increased/high expression. (D and H) Gene ontology analysis tables showing KEGG pathways, biological processes, cellular components and molecular functions that are significantly over- or under-represented in *C9orf72*-mutated (ALS) compared to isogenic microglia differentiated from either the CS52 (D) or CS29 (H) iPSC line. (I) Table showing the 20 DEGs common to CS52-C9n6-M and CS29-C9n1-M microglia. The p value and log Fold Change are indicated for each gene.

Importantly, a number of the top 30 DEGs depicted in Figure 3C (*CD14*, *CIITA*, *CSF1R*, *PIK3CG*, *Protein Tyrosine Phosphatase Receptor Type C* (*PTPRC;* also referred to as *Cluster of Differentiation 45* (*CD45*))*, Sialic Acid Binding Ig Like Lectin 1* (*SIGLEC1*; also known as *Cluster of Differentiation 169* (CD169)) encode proteins previously implicated in the modulation of pyroptosis mechanisms, including the NLRP3 inflammasome (Du et al., 2020; Lu et al., 2023; Radian et al., 2015; Souza de Lima et al., 2017; Vasudevan et al., 2022; Vidmar et al., 2019). In line with this finding, we identified genes encoding core components of the NLRP3 inflammasome, including *NLRP3*, *CASP1*, and *IL1B* among the significantly upregulated DEGs in *C9orf72*-mutated microglia, even though not among the top 30 (Supplemental Figure S1). Importantly, other genes involved in inflammasome/pyroptosis mechanisms, such as *NLRP2*, *NLR Family CARD Domain Containing 4* (*NLRC4*), *MEFV innate immunity regulator, pyrin* (*MEFV*), and *ninjurin-1* (*NINJ1*) (Barnett et al., 2023), were also among the significantly upregulated DEGs (Figure S1). These observations are consistent with the previous observation that NLRP3 activation can occur in concert with other inflammasome pathways, including NLRC4-mediated processes (Freeman et al., 2017; Ip and Medzhitov, 2015).

When the same analysis was conducted using eight RNAseq datasets from microglia derived from *C9orf72*-mutated iPSC line CS29-C9n1-M and its matching isogenic control CS29-C9n1-ISO differentiated using protocol P1, we identified 53 significantly upregulated genes in *C9orf72*- mutated microglia, while 28 genes were downregulated (Figure 3E, F). Interestingly, isogenic microglia generated from CS29-C9n1-ISO iPSCs appeared intrinsically more inflammatory than microglia generated from CS52-C9n6-ISO (Figure 2), perhaps explaining in part the lower number of DEGs compared to the latter. The list of top 30 DEGs in CS29-C9n1-M microglia revealed predominantly gene-regulation- and proliferation-associated genes, including numerous zinc finger protein-encoding genes (Figure 3G). The observation of several genes encoding zinc finger-domain proteins among the top DEGs agrees with the identification of more than 20 zinc-finger family members as genes implicated in inflammation in ALS from blood transcriptome profiles (Pappalardo et al., 2024). GO and KEGG analysis of the upregulated DEGs in CS29-C9n1-M microglia revealed enrichment in processes associated with transcription, DNA binding, RNA polymerase II activity, but no significant enrichment in immune processes apart from the term ‘response to LPS’ (Figure 3H). However, when we searched for DEGs common to both CS52-C9n6-M and CS29-C9n1-M microglia derived using protocol P1, we identified 18 significantly upregulated DEGs in both cases, as well as two common DEGs that were upregulated in CS52-C9n6-M microglia but downregulated in CS29-C9n1-M microglia (Figure 3I). These 20 common DEGs include 6 genes previously implicated in NLRP3 inflammasome-mediated mechanisms: *CD14*, *Integrin Subunit Beta 6* (*ITGB6*), *S100A9*, *S100B*, *SLAM Family Member 8* (*SLAMF8*), and *V-Set and Immunoglobulin Domain Containing 4* (*VSIG4*) (Arterbery et al., 2018; Bi et al., 2019; Huang et al., 2019; Li et al., 2025; Pruenster et al., 2023; Vizuete et al., 2022; Wang et al., 2024). This finding further suggests that mechanisms involving the NLRP3 inflammasome are dysregulated in *C9orf72*-mutated microglia.

The commonality of 20 DEGs in microglia that exhibit varying degrees of mutation-associated alterations in gene expression suggests that at least some of these transcriptomic changes may represent biologically relevant DEGs in *C9orf72*-mutated microglia. To test this possibility, and in the process search for additional DEGs, we next investigated microglia generated from CS52-C9n6-M iPSCs, and matching isogenic iPSCs, using differentiation protocol P2. As mentioned, this derivation method gives rise to microglia that appear more activated and inflammatory than those obtained with protocols 1 and 3 in our hands (Figure 2). We identified 890 genes significantly upregulated in CS52-C9n6-M microglia, with 57 genes significantly downregulated (Figure 4A, B). Perhaps not surprisingly given the phenotypic differences between microglia generated using protocols P1 and P2, the top 30 DEGs in CS52-C9n6-M microglia generated with differentiation method P2 were mainly different from the top 30 DEGs identified using differentiation method P1(Figure 4C). Importantly, however, genes associated with the NLRP3 inflammasome were also present among the top 30 DEGs identified in CS52- C9n6-M microglia generated with protocol P2. These included *Biglycan* (*BGN*), *Caveolae Associated Protein 1* (*CAVIN1*) (also termed *Polymerase 1 and Transcript Release Factor* (*PTRF*)), *Cystein Rich Angiogenic Inducer 61* (*CYR61*) (also known as *Cellular Communication Network Factor 1* (*CCN1*)), and *Periostin* (*POSTN)* (Babelova et al., 2009; Bai et al., 2010; Jin et al., 2024; Yao et al., 2020; Zheng et al., 2013; Zhou et al., 2021) (Figure 4C). Although not among the top 30 DEGs, *SIGLEC1* and *ITGB6* were also among the significantly upregulated genes in CS52-C9n6-M microglia generated with protocol P2, in agreement with protocol P1 RNAseq data. We observed that several inflammasome-associated top 30 DEGs identified using both protocols P1 and P2 encode ECM/matricellular proteins, such as *ITGB6*, *SIGLEC1*, *VSIG4*, *BGN*, *CYR61*, and *POSTN*. Upregulation of additional ECM genes was detected in *C9orf72*-mutated microglia generated with both differentiation protocols P1 and P2, including members of the non-fibrillar collagen-, laminin-, and integrin-protein families (Figure S2; Supplementary Information 1). In agreement with this, GO/KEGG analysis of molecular and cellular pathways associated with up-regulated DEGs detected using differentiation protocol 2 revealed mechanisms converging on cell adhesion, extracellular matrix organization, and cell-matrix adhesion (Figure 4D). These findings further suggest that genes involved in NLRP3 inflammasome/pyroptosis are significant DEGs in *C9orf72*-mutated ALS microglia and implicate changes at the ECM in these events.

**Figure 4.**
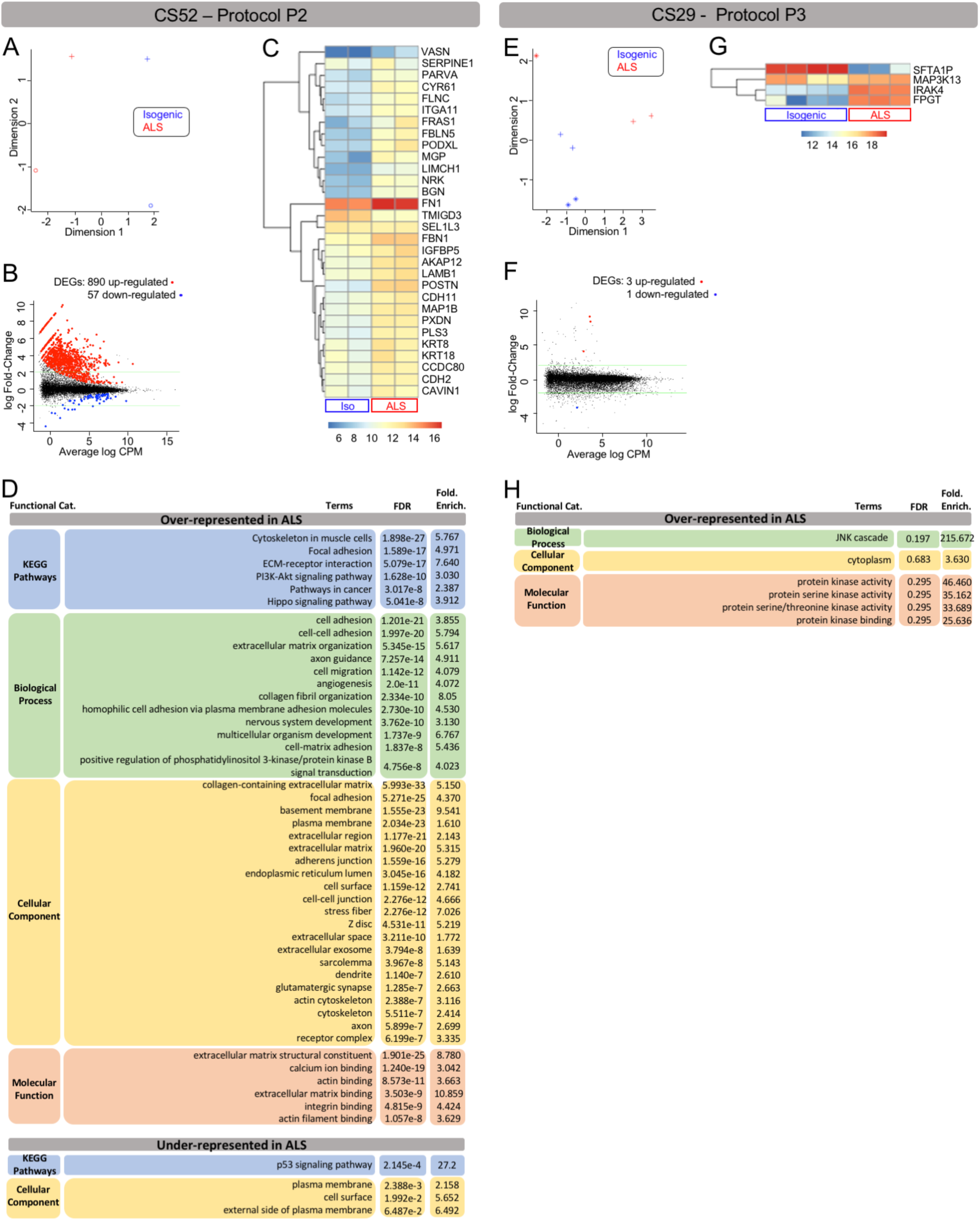
Differential gene expression between corrected isogenic and *C9orf72*-mutated microglia derived using protocols P1 and P2. (A–H) Differential gene expression and gene ontology analysis of CS52-C9n6-M versus CS52-C9n6-ISO microglia (CS52 collectively) generated using protocol P2 (A– D) or CS29-C9n1-M versus CS29-C9n1-ISO microglia (CS29 collectively) generated using protocol P3 (E–H). (A and E) Principal component analysis plots showing the relationship between the transcriptomic profiles of CS52 microglia generated with protocol P2 (A) or CS29 microglia generated with protocol P3 (E). Each plotted data point represents one biological replicate, and the different symbols refer to the different batches. Blue data points correspond to isogenic samples (Isogenic); red data points correspond to *C9orf72*-mutated samples (ALS). (B and F) Mean-difference plots (MA plots) showing the average log of Count per Million (CPM; X-axis) and log Fold Changes (Y-axis), highlighting the differentially expressed genes (DEGs) in *C9orf72*-mutated (ALS) compared to corrected isogenic microglia differentiated from the CS52 (B) or CS29 (F) iPSC lines using protocols P2 or P3, respectively. Each plotted data point represents one gene: black data points correspond to genes that are not DEG, red data points correspond to up-regulated DEGs, and blue data points correspond to down-regulated DEGs (p-adjusted <0.05, logFC >0.26). The green lines indicate 2-fold up or down. The numbers of up- and down-regulated DEGs are indicated on top of each MA plot. (C and G) Heatmaps of the topmost DEGs in *C9orf72*-mutated (ALS) compared to isogenic microglia differentiated from either the CS52 iPSC line with protocol P2 (C) or the CS29 iPSC line with protocol P3 (G). For each heatmap, each column corresponds to one sample (Isogenic or ALS, as indicated at the bottom) and each line corresponds to one gene. Relative levels of gene expression are indicated by a color scale, with cool colors corresponding to low expression and warm colors corresponding to increased/high expression. (D and H) Gene ontology analysis tables showing KEGG pathways, biological processes, cellular components and molecular functions that are significantly over- or under- represented in *C9orf72*-mutated compared to isogenic microglia differentiated from either the CS52 iPSC line with protocol P2 (D) or CS29 iPSC line with protocol P3 (H).

To further examine cell-intrinsic gene expression changes in *C9orf72*-mutated microglia, we next generated microglia from CS29-C9n1-M and CS29-C9n1-ISO iPSCs using protocol P3. As mentioned above, in our hands this method generates microglia that appear less developmentally mature and less inflammatory than those obtained using the other two methods. Although only 4 DEGs were identified in CS29-C9n1-M microglia generated using protocol P3 (Figure 4E–H), one of these DEGs, *IL-1 Receptor–Associated Kinase 4* (*IRAK4*), encodes a protein previously shown to be involved in the NLRP3 inflammasome (Leaf et al., 2017). Of note, *IRAK4* was also identified as a top 30 DEG in CS29-C9n1-M microglia generated using protocol P1 (Figure 3G). Interestingly, a second DEG was also of interest in the context of glial cell activation and caspase-mediated mechanisms of cell death, namely *Mitogen-Activated Protein Kinase Kinase Kinase 13* (*MAP3K13*, also known as *Leucine Zipper-Bearing Kinase* (*LZK*). MAP3K13/LZK promotes astrocyte reactivation and can activate NF-κB signaling (Chen et al., 2018; Masaki et al., 2003). One additional DEG was conserved in CS29-C9n1-M microglia generated using protocols P1 and P3, namely *Fucose-1- Phosphate Guanylyltransferase* (*FPGT*) (Figures 3G and 4G). *FPGT* has previously been implicated in inflammation in ALS from blood transcriptome profiles (Pappalardo et al., 2024).

In addition to the above-discussed genes, three other genes were consistently identified as top DEGs associated with the *C9orf72* mutation in all 25 RNAseq datasets, namely *Protocadherin Alpha 2* (*PCDHA2*) (FDR, 3.135e-2; fold enrichment, 4.381)*, Asparagine Synthetase Pseudogene 1* (*ASNSP1*) (FDR, 4.2e-3; fold enrichment, 6.495), and *AC004846.1* (FDR, 4.2e- 3; fold enrichment, 4.049). Of note, *AC004846.1* encodes a long non-coding RNA (lncRNA) identified as a pyroptosis-related lncRNA in colorectal cancer (Cai et al., 2022), further implicating alterations of inflammasome/pyroptosis mechanisms in *C9orf72*-mutated microglia. It is also worth mentioning that *PCDHA2*, a transmembrane protein involved in cell-cell adhesion, is a member of the *Protocadherin Alpha* gene cluster composed of 15 cadherin superfamily genes, including *PCDHA9*, which has recently been identified as a candidate gene for ALS (Zhong et al., 2024).

In summary, the combined analysis of all 25 RNAseq datasets from *C9orf72*-mutated and isogenic microglia, differentiated from different iPSC lines using different methods, identified several upregulated genes implicated in the NLRP3 inflammasome. These include genes encoding cell membrane proteins (CD14, CSFR1, PTPRC/CD45, SLAMF8), ECM components (BGN, CYR61/CCN1, ITGB6, POSTN, SIGLEC1/CD169, VSIG4), intracellular kinases (IRAK4, PIK3CG), nuclear transcription factors (CIITA), intracellular proteins secreted under inflammatory conditions (S100A9, S100B), and proteins with broad intracellular distribution (CAVIN1/PTRF). Previous studies have shown that the protein products of most of these genes cross-interact at molecular and/or cellular levels in several tissues, supporting the notion that at least some of them participate in common mechanisms in microglia (Table 1). Taken together, the present results have characterized cell-autonomous changes in the transcriptome of *C9orf72*-mutated iPSC-derived microglia, consistent across different experimental conditions, providing evidence for dysregulation of mechanisms involved in ECM biology and NLRP3 inflammasome activation.

**Table 1.** NLRP3 inflammasome-associated differentially expressed genes.

| **DEG** | **Protein Localization** | **Interaction/correlation with other DEGs** | **Involved in TLR/NF-κB signalling** |
| --- | --- | --- | --- |
| CD14 | Transmembrane | BGN, CSF1R (Roedig et al., 2019; Zheng et al., 2023) | YES (Roedig et al., 2019; Zhao et al., 2010) |
| CSF1R | Transmembrane | CD14 (Zheng et al., 2023) |  |
| PTPRC/CD45 | Transmembrane | POSTN, S110B (Rai et al., 2016; Yan et al., 2015) | YES (Puck et al., 2021) |
| SLAMF8 | Transmembrane | CD14 (Zeng et al., 2020) | YES (Qin et al., 2022; Zeng et al., 2020) |
| BGN | ECM | CD14, S100A9 (Hou et al., 2004; Roedig et al., 2019;) | YES (Roedig et al., 2019) |
| CYR61/CCN1 | ECM | CIITA, IRAK4 (Lee et al., 2010; Mooring et al., 2023) | YES (Jun and Lau, 2020) |
| ITGB6 | ECM | PIK3CG (Duggan et al., 2024) |  |
| POSTN | ECM | PTPRC (Rai et al., 2016) | YES (Zhu et al., 2024) |
| SIGLEC1/CD169 | ECM |  | YES (Li et al., 2020) |
| VSIG4 | ECM | CIITA (Kim et al., 2005) | YES (Wang et al., 2022a; Zhang et al., 2021) |
| IRAK4 | Intracellular | CD14, CYR61 (Mooring et al., 2023; Sayers et al., 2024) | YES (Kim et al., 2007; Kim et al., 2019) |
| PIK3CG | Intracellular | ITGB6 (Duggan et al., 2024) |  |
| S100A9 | Intracell/Secreted | BGN (Hou et al., 2004) | YES (Hermani et al., 2006; Riva et al., 2012) |
| S100B | Intracell/Secreted | PTPRC (Yan et al., 2015) | YES (Bianchi et al., 2010; Villarreal et al., 2011) |
| CIITA | Nucleus | CYR61, VSIG4 (Kim et al., 2005; Lee et al., 2010) | YES (Benson & Ernst, 2009; Qian et al., 2018) |
| CAVIN1/PTRF | Nucleus, caveolae |  | YES (Yi et al., 2022; Zheng et al., 2013) |

Shown are the localizations of the encoded proteins (ECM, extracellular matrix), molecular interactions and/or expression correlations among different DEGs, and involvement in mechanisms regulating TLR and NF-κB signalling.

### Effect of C9orf72-mutated iPSC-derived microglia conditioned medium on motor neurons

The cell-autonomous transcriptomic changes observed in *C9orf72*-mutated microglia suggest that these cells can be toxic to motor neurons at least in part due to dysregulated inflammasome-activated mechanisms leading to release of neurotoxic factors. To determine whether *C9orf72*- mutated microglia secrete molecules that are sufficient to decrease motor neuron survival, we examined the effect of conditioned media from either *C9orf72*-mutated microglia or their matching isogenic controls on both *C9orf72*-mutated and isogenic motor neurons. Mutated or isogenic motor neurons were differentiated for 28 days, followed by addition of either no conditioned media or conditioned media collected from *C9orf72*-mutated or isogenic microglia generated using protocol P1. Motor neurons were then cultured till day 49 and then subjected to immunocytochemistry with an antibody against cleaved caspase 3 (CC3) to detect apoptotic cells (Figure 5A). Motor neurons were visualized with an antibody against choline acetyltransferase (CHAT). With no conditioned media added, quantification of the numbers of CHAT-expressing cells positive for CC3 showed increased numbers of apoptotic *C9orf72*- mutated motor neurons compared to isogenic motor neurons (Figure 5B, bars 1 and 2) indicating intrinsic reduced viability of the mutant cells, in agreement with previous studies (Thiry et al., 2024). Exposure to conditioned media from isogenic microglia had no detectable effect on the viability of either *C9orf72*-mutated or isogenic motor neurons (Figure 5B, bars 1– 4). In contrast, we observed a statistically significant increase in the fraction of CC3-positive *C9orf72*-mutated motor neurons in the presence of conditioned media from *C9orf72*-mutated microglia compared to absence of any conditioned media or exposure to media from isogenic microglia (Figure 5B, cf. bars 2, 4, 6). Isogenic motor neuron cultures tended to contain more CC3-positive cells in the presence of conditioned media from *C9orf72*-mutated microglia, but this was not statistically significant when compared to conditioned media from isogenic microglia (Figure 5B, cf. bars 3 and 5). These results provide evidence that *C9orf72*-mutated microglia enhance the death of *C9orf72*-mutated motor neurons, and that this effect does not require motor neuron-microglial cell contact.

**Figure 5.**
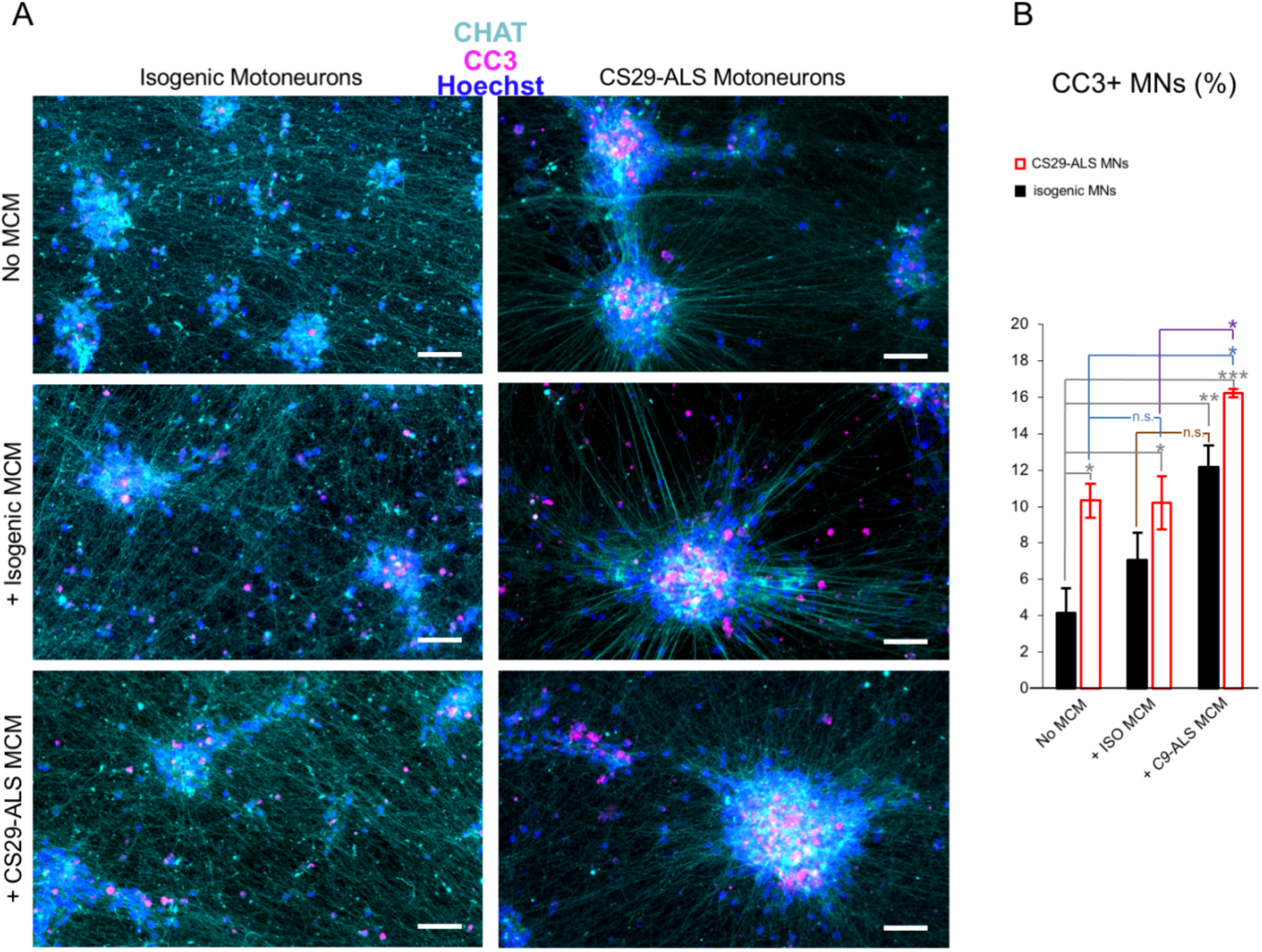
Effect of *C9orf72*-mutated microglia conditioned media on *C9orf72*-mutated motor neurons. (A) Representative immunocytochemistry images of motor neurons (Motoneurons) derived from isogenic (left column) or *C9orf72*-mutated (CS29-ALS) (right column) iPSCs and maintained in culture for 3 weeks in the absence of microglia conditioned media (MCM) (top images: No MCM), or in the presence of conditioned media from isogenic microglia (middle images: isogenic MCM) or *C9orf72*-mutated (CS29-ALS) MCM (bottom images). Cultures were immunostained with anti-CC3 antibody (magenta), anti-CHAT antibody (cyan), and counterstained with Hoechst (blue). Scale bars, 50 µm. (B) Bar plot showing the quantification of the percentage of apoptotic CC3-positive motor neurons in isogenic or *C9orf72*-mutated CS29 motor neuron preparations after 3 weeks of culture in each of the following three conditions: absence of MCM (No MCM); presence of isogenic microglia conditioned media (ISO MCM); or presence of *C9orf72*-mutated CS29 MCM (C9-ALS MCM). Statistics: two-way ANOVA and Tukey’s multiple comparisons post-test; n.s. = non-significant; * = p value < 0.05; ** = p value < 0.005; *** = p value < 0.0005.

## Discussion

There is evidence from studies in animal and human cell models of ALS that the *C9orf72* HRE is associated with activation of the NLRP3 inflammasome in microglia (Banerjee et al., 2023; Fu et al., 2023; Rivers-Auty et al., 2024; Trageser et al., 2023). Little information is available, however, about the molecular mechanisms underlying the activation of inflammasome-mediated processes in *C9orf72*-mutated microglia. It is hypothesized that a variety of microglia-intrinsic (e.g., homeostasis-altering events) and/or microglia-extrinsic (e.g., host damage-associated molecular patterns) mechanisms can activate inflammasome PRRs in *C9orf72*-HRE microglia. These PRRs include, but are not limited to, TLR and TNF receptor family members (Barnett et al., 2023; Fu and Wu, 2023; Sharma and Kanneganti, 2021). In this study, we sought to identify cell-intrinsic changes in the transcriptome of *C9orf72*-HRE microglia that might shed light on the molecular events underlying inflammasome activation in ALS/FTD microglia.

Our investigations have identified several genes upregulated in *C9orf72*-mutated microglia. Elevated expression of a number of proteins encoded by these genes could cause dysregulation of TLR/NF-κB-mediated activation of the NLRP3 inflammasome. For instance, DEG *SLAMF8* codes for a cell surface receptor that is highly induced in collagen-induced rheumatoid arthritis and knockout of *SLAMF8* attenuates this condition through inhibition of TLR4/NF-κB signaling (Qin et al., 2022). Consistently, simultaneous knockout of *SLAMF8* and its related family member *SLAMF9* prevents LPS-induced liver inflammation by downregulating TLR4 expression on macrophages (Zeng et al., 2020). These findings suggest that abnormally elevated levels of SLAMF8 in *C9orf72*-mutated microglia might play a role in initiating inflammation and/or maintaining a pro-inflammatory environment via TLR4/NF-κB-mediated activation of the NLRP3 inflammasome. Another candidate activator of the NLRP3 inflammasome is the product of DEG *CD14*, which is a GPI-anchored co-receptor for TLRs involved LPS-induced responses. CD14/TLR coreceptors are important for microglia activation in ALS, as demonstrated by the finding that microglia activation mediated by extracellular mutant superoxide dismutase 1 was attenuated using TLR2, TLR4 and CD14 blocking antibodies, or when microglia lacked CD14 expression (Zhao et al., 2010). Thus, elevated levels of CD14 in *C9orf72*-mutated microglia could cause increased TLR signaling, with the potential to activate the NLRP3 inflammasome via NF-κB. A role for CD14 in dysregulated TLR activation in *C9orf72*-mutated microglia is also suggested by the present identification of *BGN* as an upregulated gene in these cells. The *BGN* gene product, biglycan (BGN), is an ECM-derived damage-associated molecular pattern that acts as a high affinity ligand for CD14 in macrophages, where CD14 is required for BGN-mediated TLR2/4-dependent inflammatory signaling pathways (Roedig et al., 2019). Consistently, BGN can activate the NLRP3 inflammasome in macrophages leading to activation of CASP1 and release of IL1B (Babelova et al., 2009). These observations suggest that elevated levels of both BGN and CD14 cause dysregulated NLRP3 inflammasome activation through elevated TLR/NF-KB signaling.

The present study has identified other ECM protein-encoding genes, in addition to *BGN*, among DEGs in *C9orf72*-mutated microglia with the potential to activate TLR/NF-κB-mediated mechanisms. More specifically, the matricellular protein CYR61/CCN1, encoded by DEG *CYR61*, acts as a pattern recognition receptor that binds bacterial pathogen-associated molecular patterns in macrophages, where it interacts directly with TLR2 and TLR4 to activate inflammatory responses (Jun and Lau, 2020). In agreement with this finding, CYR61/CCN1 promotes the expression of NLRP3, ASC, CASP1, and IL1B in astrocytes in an autocrine manner (Jin et al., 2024). Separate investigations showed that knockdown of the DEG *POSTN*, encoding periostin (POSTN), a secreted matricellular protein, suppresses CASP1-mediated pyroptosis and NLRP3 inflammasome activation in muscle cells (Yao et al., 2020). Consistently, POSTN activates inflammation in atopic dermatitis and modulates the immune response and ECM remodeling during atherosclerotic plaque formation (Nunomura et al., 2023; Schwanekamp et al., 2016). Relevant to these observations, POSTN enhances integrin/NF-κB signaling during the process of intervertebral disc degeneration (Zhu et al., 2024). Although not among the top 30 DEGs found in our datasets, numerous additional ECM genes were present among the statistically significant genes dysregulated in *C9orf72*-mutated microglia, consistent with alterations in ECM composition. Together, these observations suggest that the *C9orf72* HRE causes dysregulation of mechanisms that may underlie an interplay between ECM biology and inflammatory responses in microglia. This notion is consistent with the previous observation of a link between NLRP3 inflammasome/pyroptosis and ECM remodelling in non-neural systems (He et al., 2023), further suggesting a connection between dysregulated inflammasome activation and the ECM in *C9orf72*-mutated ALS microglia.

Perturbations of interplays between the ECM and inflammatory mechanisms is expected to affect several microglial cell functions, including cell adhesion and cell migration. In this regard, DEG *SIGLEC1* encodes a cell adhesion molecule that binds sialylated ligands on endogenous or pathogenic membrane surfaces (Herzog et al, 2022). Inhibition of *SIGLEC1* expression in monocytes reduces levels of proinflammatory cytokines such as IL1B, IL6, and IL8 (Wang et al., 2022b). Moreover, SIGLEC1 exerts a proinflammatory effect in chronic obstructive pulmonary disease through activation of NF-KB (Li et al., 2020). These observations identify SIGLEC1 as another cell surface protein potentially involved in pro-inflammatory processes in microglia.

Our results also suggest intracellular mechanisms by which the NLRP3 inflammasome may be stimulated through dysregulated activation of TLR/NF-κB signaling. Interleukin 1 Receptor Associated Kinase 4, encoded by DEG *IRAK4*, is a kinase that activates NF-κB downstream of TLR signaling. Mice lacking *IRAK4* display severe impairment of TLR signaling because of the loss of the normal physical association of the cytoplasmic portion of TLRs with IRAK4 (Suzuki et al., 2002). Consistently, inhibition of IRAK4/NF-κB/NLRP3 pathway reduces pyroptosis in hippocampal neurons (Zhao et al. 2024). Thus, it is reasonable to speculate that upregulation of IRAK4 in *C9orf72*-mutated microglia is one of the events contributing to elevated NLRP3 inflammasome, especially if occurring together with other dysregulated mechanisms leading to TLR activation. Another potential mechanism by which the NLRP3 inflammasome could become activated in *C9orf72*-mutated microglia through enhanced TLR signaling is suggested by the present identification of *S100A9* and *S100B* as upregulated genes. Both these genes encode intracellular proteins containing EF-hand calcium-binding motifs. During inflammation, S100A9 and S100B proteins are released in the extracellular milieu where they can bind to TLRs and Receptor for Advanced Glycation End-products (RAGE) (Franz et al., 2022; Villareal et al., 2014). In microglia, increased level of extracellular S100B upregulates IL1B and TNF-a expression via RAGE engagement and this effect requires activation of NF-κB (Bianchi et al., 2010). Extracellular S100B can promote reactive astrocyte transformation leading to a pro-inflammatory phenotype characterized by expression of TLR2 and IL1B (Villareal et al., 2014). Consistently, extracellular S100A9 stimulates macrophages to undergo NLRP3 inflammasome activation (Sheng et al., 2024). Conversely, ablation of S100A9 in macrophages dampens NLRP3-driven inflammatory response and pyroptosis (Sheng et al., 2023). Of note, *S100A9* expression is downregulated when NLRP3 inflammasome blockade leads to reduced inflammation and ECM remodeling in adipose tissue (Unamuno et al., 2021). Taken together, these observations suggest several mechanisms by which the upregulated genes identified in *C9orf72*-mutated microglia in this study could be involved in NLRP3 inflammasome activation.

The intrinsic inflammatory phenotype of *C9orf72*-mutated microglia suggests that these cells are toxic to motor neurons. This possibility agrees with the results of previous studies showing that LPS-primed *C9orf72*-mutated iPSC-derived microglia promote the death of healthy motor neurons in microglia-motor neuron co-culture studies (Banerjee et al., 2023; Vahsen et al., 2023). We examined the possibility that factors released by *C9orf72*-mutated microglia are sufficient to decrease the survival of *C9orf72*-mutated motor neurons in the absence of cell-cell contact. The present studies have shown that exposure to conditioned media from *C9orf72*- mutated microglia, but not isogenic microglia, increases the number of apoptotic *C9orf72*- mutated motor neurons. The effect of the *C9orf72*-mutated microglia conditioned media on isogenic motor neurons was only moderate. This finding provides evidence that *C9orf72*- mutated microglia are primed to be deleterious to motor neurons in a cell-autonomous fashion and that this effect is mediated by factors released by microglia. At least some of the proteins encoded by the *C9orf72*-mutated microglia DEGs identified in this study could contribute to mechanisms of motor neuron degeneration. Both S100A9 and S100B, when secreted by glial cells, can mediate neurotoxic effects. Elevated levels of extracellular S100B are involved in the death of enteric neurons caused by 5-fluorouracil in a RAGE/NF-κB-dependent pathway (Costa et al., 2019). Moreover, acute exposure to extracellular S100B alters the electrophysiological activity of tyrosine hydroxylase-expressing dopaminergic neurons in primary mouse midbrain cultures and S100B overexpression in mice accelerates the loss of substantia nigra pars compacta dopaminergic neurons (Bancroft et al., 2022). S100A9 is pro-inflammatory and highly amyloidogenic in the amyloid-neuroinflammatory cascade in neurodegenerative diseases leading to cognitive impairment (Andrade-Talavera et al., 2022). Consistently, S100A9 inhibition in microglia and macrophages by pharmacological treatment protects against neuronal death *in vitro* and brain injuries *in vivo* in a model of ischemic stroke (Liu et al, 2024).

In conclusion, the transcriptomic changes in *C9orf72*-mutated microglia described in this study provide evidence for a cell-autonomous pro-inflammatory phenotype associated with this mutation. Moreover, they agree with earlier evidence of NLRP3 inflammasome activation in *C9orf72*-mutated microglia and provide previously unavailable information on the identity of several genes that could contribute to inflammasome activation. Some of the proteins encoded by these differentially expressed genes have the potential to participate in both cell-autonomous mechanisms of microglia-mediated inflammation and non-cell autonomous processes of microglia-induced motor neuron degeneration in ALS/FTD. At least some of the mechanisms mediated by these gene products could provide targets for translational research.

## Materials and Methods

### Human induced pluripotent stem cells

Four previously studied (Chi et al., 2023; Ho et al., 2021; Gao et al., 2024; Thiry et al., 2024) human iPSC lines were used for microglia derivation: the *C9orf72*-mutated iPSC lines CS52iALS-C9n6 (CS52-C9n6-M) and CS29iALS-C9n1 (CS29-C9n1-M), and their matching isogenic control lines CS52iALS-C9n6.ISOxx (CS52-C9n6-ISO) and CS29iALS-C9n1.ISOxx (CS29-C9n1-ISO). These iPSC lines were obtained from Cedars-Sinai (Los Angeles, CA, USA). Undifferentiated state of human iPSCs was assessed by testing for expression of the stem cell markers NANOG and OCT4 using rabbit anti-NANOG (1/1,000; Abcam; Cambridge, UK; Cat. No. ab21624) and rabbit anti-OCT4 (1 μg/ml; Abcam; Cat. No. ab19857) or goat anti- OCT3/4 (1/500; Santa Cruz Biotechnology; Dallas, TX, USA, Cat. No. sc-8628) antibodies, as previously described (Chen et al., 2021).

### Derivation of microglia from human iPSCs

Derivation of microglia from human iPSCs was performed essentially as described in three separate protocols published by Fossati and coworkers (Douvaras et al., 2017), Blurton-Jones and colleagues (McQuade et al., 2018), and Cowley and colleagues (Haenseler et al. 2017). Since we used slightly modified versions of these methods, they are referred to as protocols “P1”, “P2” and “P3”, respectively. The modifications made to the Douvaras et al. and McQuade et al. methods were previously described (Tang et al., 2022). The following modifications were made to the Haenseler et al. method. Floating progenitor cells were harvested from embryoid bodies weekly and cultured by plating onto 6-well plates or glass coverslips coated with Corning® Matrigel (MilliporeSigma; Burlington, MA, USA; Cat. No. 354277) at 100,000 per cm^2^. Cells were cultured for 10-20 days in DMEM/F12 (Thermo-Fisher Scientific; Waltham, MA, USA; Cat. No. 10565-018) with 100 ng/mL IL-34 (Thermo-Fisher Scientific; Cat. No. 200-34), 10 ng/mL GM-CSF (R&D Systems; Minneapolis, MN, USA; Cat. No. 215-GM), N2 supplement (Thermo-Fisher Scientific; Cat. No. 17502-048), 2 mM GlutaMAX (Thermo- Fisher Scientific; Cat. No. 35050-061), and antibiotic-antimycotic (Thermo-Fisher Scientific; Cat. No. 15240-062). All microglia preparations generated using these three protocols were validated through morphological analysis and immunocytochemistry as described (Tang et al., 2022) before being subjected to RNAseq.

### RNA sequencing

Total RNA was isolated for RNAseq from cell pellets obtained from microglia generated with each one of the three protocols described above using sequential treatment with TRIzol Reagent (ThermoFisher Scientific; Cat. No. 15596026) and PureLink RNA Micro Scale Kit (ThermoFisher Scientific; Cat. No. 12183-016) following the instructions provided by the manufacturer. We isolated RNA from iPSC-derived microglia at a series of time points spanning days *in vitro* 45 to 66 for preparations generated with protocols P1 and P2 and days *in vitro* 75 to 108 for preparations induced with protocol P3. These time points were long enough for induced cells to acquire a microglia phenotype according to all three derivation methods (Douvaras et al., 2017; Haenseler et al., 2017; McQuade et al., 2018). All RNA samples were analyzed by Illumina next-generation sequencing at the Genomics platform at the Institute for Research in Immunology and Cancer, Montreal, Quebec, Canada (https://www.iric.ca/en/research/platforms-andinfrastructures/genomics). Adaptor sequences and low-quality bases in the resulting FASTQ files were trimmed using Trimmomatic version 0·35 (Bolger et al., 2014), and genome alignments were conducted using STAR version 2·5 1b (Dobin et al., 2013). Sequences were aligned to the human genome version GRCh38, with gene annotations from Gencode v29 based on Ensemble release 94. As part of quality control, the sequences were aligned to several different genomes to verify that there was no sample contamination. Raw read-counts were obtained directly from STAR and reads in transcripts per million (TPM) formats were computed using RSEM (Li & Dewey, 2011).

### In silico analysis

To compare isogenic microglia preparations derived with three different protocols, heatmaps showing the levels of expression of 49 microglia/macrophage marker genes (Butovsky et al., 2014; Hickman et al., 2013; Muffat et al., 2016; Zhang et al., 2014) were generated for each sample. Read counts for these specific genes were converted into log2-counts-per-million (logCPM) values as previously described (Tang et al., 2022). Accordingly, negative values represent very low gene expression values. Hierarchical clustering of rows was applied to the heatmaps to group genes with correlated expression patterns, resulting in different gene lists, depending on the samples analyzed in the given heatmap. For each heatmap, color comparisons can be made across samples for each gene as well as between genes for any given sample.

Differential gene expression analysis was conducted on the raw read-count matrix using an edgeR (version 4·0) pipeline as previously described (Chen et al., 2016). Briefly, genes were filtered to retain only those with a robust expression level. A scaling factor of the trimmed mean of M-values (TMM) between each pair of samples was used to normalize the differing RNA composition of each sample. The differentiation protocol and experimental batch were incorporated into the design used for differential expression analysis since they were the two known sources of variability in the studied samples. The “glmQLFit” and “glmQLFTest” functions of edgeR were then used to obtain DEGs with the quasi-likelihood (QL) method. Genes were considered statistically significant if the adjusted p-value (false discovery rate; FDR) was <0.05, with no fold-change cutoff applied to restrict the number of DEGs identified.

Gene set enrichment analysis (GSEA) was conducted on the DEGs using the Functional Annotation Tool of DAVID Bioinformatics Resources 6·8 (Huang da et al., 2009a; 2009b). GO terms and KEGG pathways enriched in the DEGs were computed separately for upregulated and downregulated genes, as this increases the statistical power to identify pathways/terms pertinent to the phenotype of interest (Hong et al., 2013). For each GO category (e.g., Biological Process or Molecular Function), the “direct” function in DAVID was used to restrict the results to those directly annotated by the source database and to avoid repetitive and generic parent terms. For the KEGG pathways, only the non-disease-specific pathways are shown. For both GO and KEGG pathways analysis, all terms with associated FRD < 1 are shown, except for categories presenting more than 10 terms, in which case only the more statistically significant terms with associated FDR<e^-6^ are shown.

### Open-source data

Raw read counts from RNA sequencing of undifferentiated human iPSCs were obtained from the LINCS Data Portal (LSC-1002 and LSC-1004 from Dataset LDS-1355) (http://lincsportal.ccs.miami.edu/datasets/). LSC-1002 and LSC-1004 correspond to human iPSC lines CS14i-CTR-n6 and CS25iCTR-18n2, provided by Cedar Sinai Stem Cell Core Laboratory. These raw data were combined with the RNAseq data from the present microglia preparations and analyzed using a single pipeline, to generate heatmaps comparing the levels of expression of activated microglia marker genes in each sample.

### 4.6. Incubation of cultures of iPSC-derived motor neurons with microglia conditioned media

Motor neuron cultures enriched in phrenic motor neurons were derived from human iPSC lines CS29-C9n1-M and its matching isogenic control line CS29-C9n1-ISO according to a previously described protocol (Thiry et al., 2024). Briefly, neural progenitor cells were dissociated on day 6 and split 1:5 with a chemically defined neural medium including DMEM/F12 supplemented with GlutaMAX (1/1; Thermo-Fisher Scientific; Cat. No. 35050-061), Neurobasal medium (1/1; Thermo-Fisher Scientific; Cat. No. 21103-049), N2 (0.5X; Thermo-Fisher Scientific; Cat. No. 17504-044), B27 (0.5X; Thermo-Fisher Scientific; Cat. No. 17502-048), ascorbic acid (100 μM; Sigma-Aldrich; Oakville, ON, Canada; Cat. No. A5960), antibiotic-antimycotic (1X; Thermo-Fisher Scientific; Cat. No. 15240-062), supplemented with retinoic acid (RA) (1 µM, Sigma-Aldrich; Cat. No. R2625) and purmorphamine (0.125 µM, Sigma-Aldrich; Cat. No. SML-0868) in combination with 1 μM CHIR99021 (STEMCELL Technologies; Vancouver, BC, Canada; Cat. No. 72054), 2 μM DMH1 (Sigma-Aldrich; Cat. No. D8946) and 2 μM SB431542 (Tocris Bioscience; Bristol, UK; Cat. No. 1614). The culture medium was changed every other day for 6 days and the resulting motor neuron progenitors were then expanded for 6 days with the same medium containing 3 µM CHIR99021, 2 µM DMH1, 2 µM SB431542, 1 µM RA, 0.125 µM purmorphamine and 500 µM valproic acid (Sigma-Aldrich; Cat. No. P4543). Motor neuron progenitors were dissociated and plated at 50,000 cells/well on coverslips coated with Dendritic Polyglycerol Amine/Matrigel substrate as described previously (Thiry et al., 2022). Motor neurons were differentiated in the same neural medium supplemented with 1 µM RA and 0.125 µM purmorphamine, 0.1 µM Compound E (MilliporeSigma; Cat. No. 565790), insulin-like growth factor 1 (10 ng/mL; R&D Systems; Cat. No. 291-G1-200), brain-derived neurotrophic factor (10 ng/mL; Thermo-Fisher Scientific; Cat. No. PHC7074) and ciliary neurotrophic factor (10 ng/mL; R&D Systems; Cat. No. 257-NT-050). Motor neuron culture medium was replaced every other day for one week. Motor neuron cultures were validated by immunocytochemistry and single-cell RNA sequencing as described (Thiry et al., 2020; 2022; 2024). For studies with microglia conditioned media, after one week in motor neuron culture medium, the medium was replaced every other day for an additional 2-weeks with a mixture of motor neuron culture medium (50%) and microglia conditioned medium (50%) obtained from either *C9orf72*-mutated or isogenic microglia derived from CS29-C9n1-M or CS29-C9n1-ISO iPSCs using protocol P2, as described above. After 46 days of culture (3-weeks post-plating), culture media were removed abd motor neurons were rinsed and analyzed by immunocytochemistry.

### Analysis of apoptosis in iPSC-derived motor neuron cultures

Immunocytochemistry with rabbit anti-Cleaved-Caspase 3 (CC3 (Asp175); 1/400; Cell Signaling; New England Biolabs, Ltd., Ontario, Canada; Cat. No. 9661S) in combination with goat anti-CHAT antibody (1/100; Millipore; Cat. No. MAB144P) was performed to detect apoptotic motor neurons in cultures obtained from CS29-C9n1-M or CS29-C9n1-ISO iPSCs, in the absence or presence of conditioned media from CS29-C9n1-M or CS29-C9n1-ISO microglia. For quantification, images in 3 random fields were taken with a 20X objective using an Axio Observer Z1 microscope connected to an AxioCam camera using ZEN software (Zeiss) and analyzed with Image J.

### Statistical analysis of motor neuron cell death assays

To account for culture variability and ensure experimental reproducibility, studies with motor neurons plus or minus microglia conditioned media were performed with at least 3 biologically independent cultures per condition. Error bars shown are means ± standard error of means (SEM) of the average. Two-way repeated measure ANOVA and Tukey’s multiple comparisons post-hoc test were used to detect differences in the proportion of CC3-positive motor neurons in isogenic and *C9orf72*-mutated-derived cells in each of the three culture conditions tested, namely absence of microglia conditioned media, presence of isogenic microglia conditioned media, or presence of *C9orf72*-mutated microglia conditioned media.

## Data Availability

R code used for in silico analysis was based on the published code (Tang et al., 2022). The raw data are available at the Sequence Read Archive (SRA): http://www.ncbi.nlm.nih.gov/bioproject/1235672.

## Acknowledgments

We thank Rita Lo for excellent lab support and Thomas Durcan for providing access to scientific equipment. These studies were funded in part by the Canadian Institutes for Health Research and Fonds de la recherche en Santé-Quebec under the frame of E-Rare-3, the ERA-Net for Research on Rare Diseases (SS) and by ALS Canada (SS). S.S is a Distinguished James McGill Professor of McGill University.

## Author contributions

NS Pulimood: conceptualization, methodology, investigation, analysis and interpretation.

L Thiry: conceptualization, methodology, investigation, analysis and interpretation, writing.

Y Tang: methodology, investigation, analysis and interpretation.

S Stifani: conceptualization, methodology, analysis and interpretation, resources, funding acquisition, supervision, writing.

## Supplementary Figures and Information

**Figure S1.**
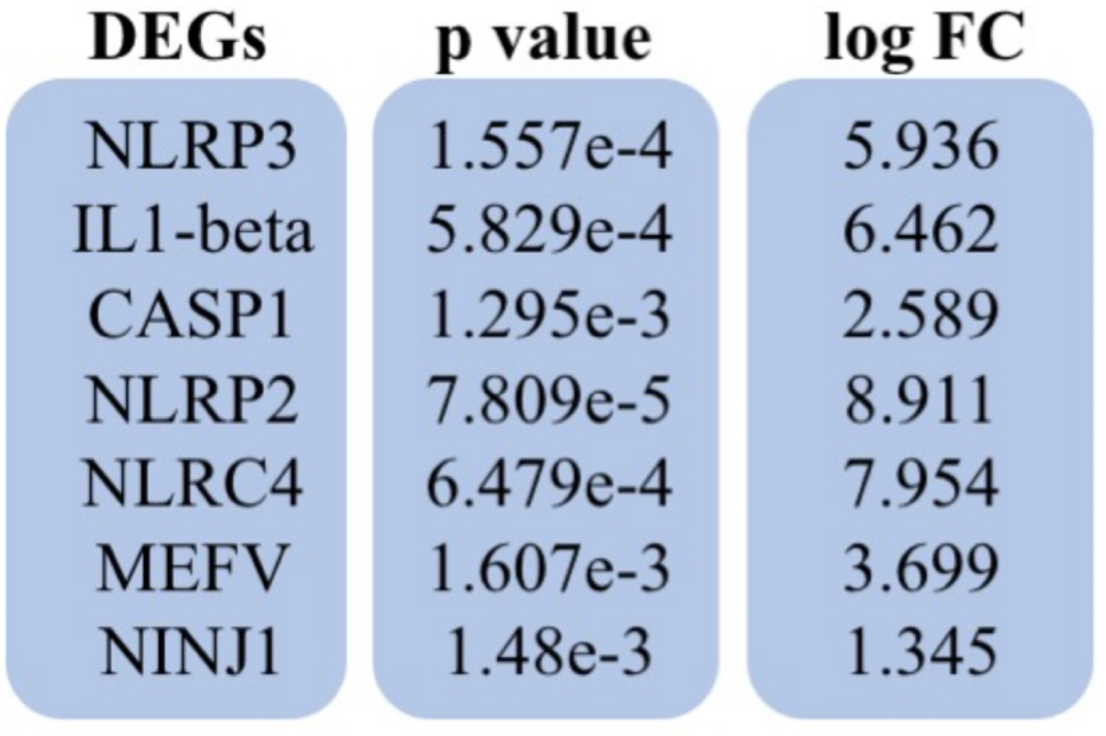
Dysregulation of inflammasome genes in *C9orf72*-mutated microglia generated using derivation protocol P1. Shown are the p value and log Fold Change (logFC) for the indicated DEGs in CS52-C9n6-M versus CS52-C9n6-ISO microglia.

**Figure S2.**
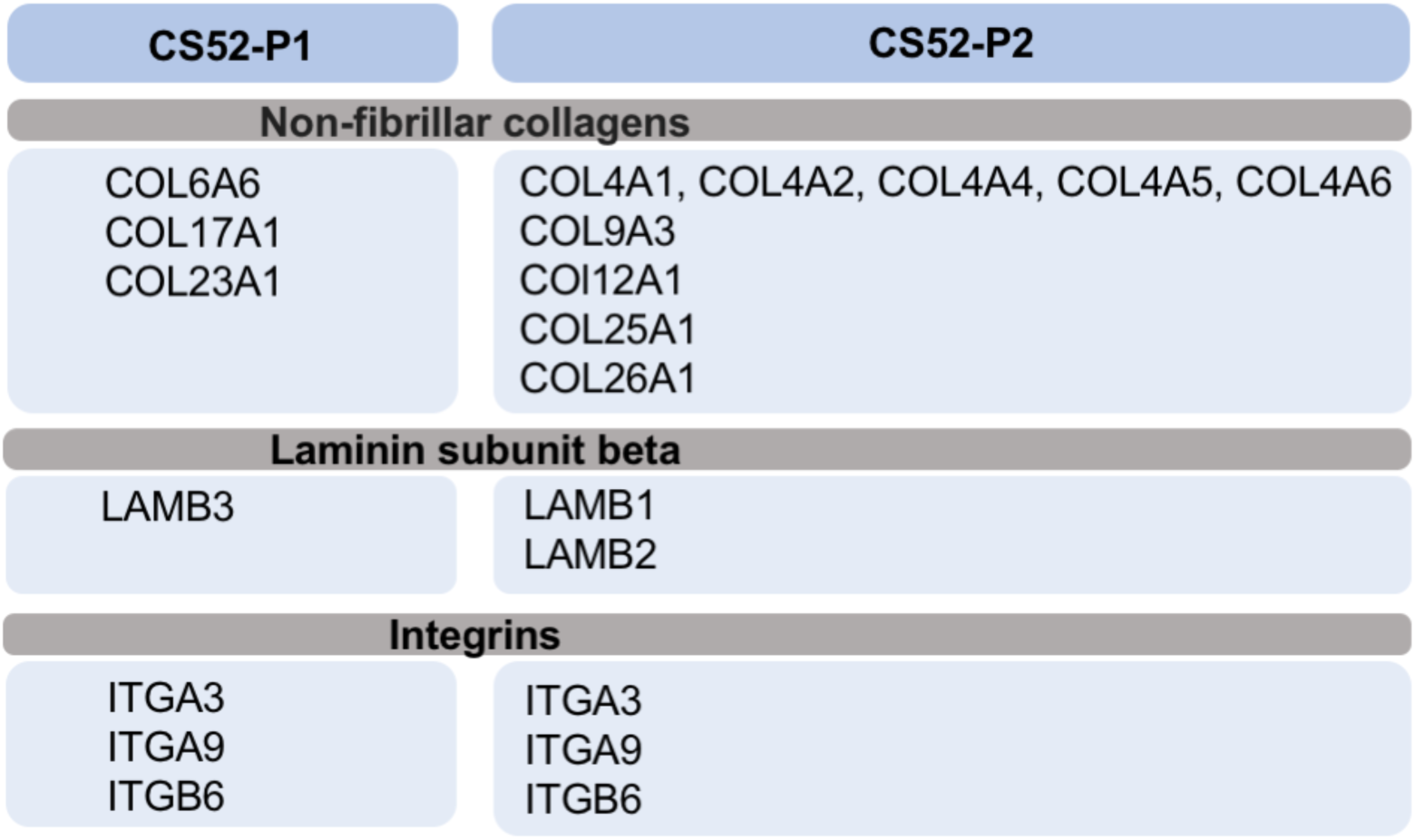
Dysregulation of extracellular matrix genes in *C9orf72*-mutated microglia generated using derivation protocols P1 and P2. Shown are genes encoding members of the non-fibrillar collagen, laminin subunit beta, and integrin families dysregulated in microglia generated using both differentiation protocols P1 and P2. CS52 = CS52-C9n6-M versus CS52-C9n6-ISO microglia; P1 and P2 = differentiation protocols P1 and P2.

## Protocol 1: 614 Up-regulated DEGs in CS52-C9n6-M microglia

FGR, LAP3, CASP10, HSPB6, SKAP2, ITGAL, TRAF3IP3, STAB1, CD4, BTK, TYROBP, GABRA3, CD22, WAS, SLC11A1, CD74, RUNX3, NLRP2, SLC45A4, SLAMF7, TNFRSF1B, STAP1, MSR1, CAPG, LCP2, COL23A1, KCNQ1, CBLN4, PRDM1, TBXAS1, PARP12, GNA15, EYA2, ANKRD44, COL17A1, CTNNA2, CD84, SPI1, TRIP13, PTPN18, FCGR2B, SIDT1, DHRS9, ST6GAL1, SCARF1, CTTNBP2, DCX, MAP2, ACER3, MKNK1, RBL1, MEF2C, PTPRC, FYB1, EPB41L3, GRHL2, SMAP2, PILRA, NME8, SNX10, SIGLEC1, OAS1, ARHGAP4, LYZ, CD209, NLRC4, TBX15, ADA2, GABRP, BLNK, TREM2, NRP1, CYTH4, MFNG, GRK3, CRYBB1, AP1B1, HMOX1, PDGFB, APOL4, NCF4, CSF2RB, LGMN, CEP128, MMP9, ABHD12, CD40, CTSZ, TUBB1, RASSF2, SIRPB1, HCK, SAMHD1, PPP1R16B, TLR8, RENBP, SMS, ACP5, CORO1A, MEFV, ATP8B4, BMF, DMXL2, FGL1, CD37, LYL1, IL4I1, RASAL3, MYH14, CD33, LILRB5, ARRDC2, PIK3CG, TFEC, CHCHD2, GIMAP2, TRIM14, DOCK8, APBA1, RASSF4, CXCL12, PKD2L1, VSIR, LIPA, ABI3, FAM20A, SLC2A9, CTSC, UNC93B1, MS4A6A, MS4A4A, IL10RA, SLC15A3, SELPLG, BIN2, OAS3, OAS2, ARHGDIB, PTPN6, FRK, CRYBG1, LY86, RNF130, PCDH12, CD86, FOXP1, TFCP2L1, CYTIP, ITGB6, GALNT3, CCDC88A, KYNU, PLEK, NCF2, CD48, CTSD, MFSD1, CNTRL, FLVCR2, MOB3B, CCDC170, CXorf21, KCNJ5, ADCY7, CD80, CAT, CCRL2, TNFSF10, TMIGD3, SASH3, HS3ST2, SRGN, HVCN1, ADGRE5, ARHGAP9, NCKAP1L, SH3TC1, IL1B, ATG4C, CD93, EVI2A, FGD3, ADGRE2, TUBA4A, YWHAH, DOCK4, LRRC4, IRF5, WDFY4, TMOD2, CD68, SIGLEC9, RNASE1, STARD8, CNN1, LSP1, TNNI2, GMFG, THEMIS2, MPP1, LILRB2, NINJ1, ZNF132, EMILIN2, CARD6, CLEC10A, XAF1, ALOX5AP, CHIT1, PRAM1, CNDP2, GIMAP6, GIMAP4, AMPD3, HSD17B4, ADAMDEC1, CD180, CAMK1, SYT6, VAV3, PTPN22, DOCK2, SLC43A3, C5AR2, SLC37A2, HAVCR2, SDS, DTX1, CD36, LMO2, PRRG4, MICAL1, TEC, CPM, NPL, DOCK10, RCBTB2, AOAH, MYO1G, TM6SF1, ERMN, IL10, IL1RN, TLR4, IRF4, FLOT1, RNF144B, TLR2, MYO7A, SLCO2B1, CASP1, KNL1, PLCB2, TMEM62, ZNF280D, IFI44L, IFI44, CALHM2, IDH1, MNS1, CENPE, PARVG, PPFIA2, LGR5, GAS2L3, ACVRL1, GPR84, BRCA2, ITGB7, LPAR6, VPS37B, RTN1, SLC38A6, GPR65, PSTPIP1, ZNF710, ITGAX, IGSF6, IRF8, ARRB2, PIK3R5, TNFRSF11A, VAV1, TRPM2, MYO1F, SIGLEC10, RCN3, CD53, FCGR2A, RGL1, S100A8, EPHX1, ITGA9, STAC, ADPRH, NCEH1, TCTA, SCD5, OTULINL, IQGAP2, PLA2G7, DOK3, FGD2, ARHGAP18, SLC18B1, ARMT1, TRIM4, TMEM140, SH3KBP1, DOK2, ST18, GINS4, ATP6V0D2, TMEM236, LRMDA, HTR7, PTPRJ, ST14, FERMT3, FCGR1A, TMEM86A, PTPRO, TDO2, KCNE4, ARL11, PDE3B, KCNK13, RASGRP3, ANKRD22, MR1, GRAP, FGD5, PLCL2, SAMSN1, PIK3AP1, VSIG4, SLA, SLC28A1, TDRD9, MAPK13, TSPAN18, TNFRSF14, TSPAN33, SLAMF8, FCER1G, C1QC, ACE, ITGB2, S100B, JAML, CCR5, HK3, ZNF385A, SCIMP, TNFSF13, TAL1, LAPTM5, NLRP3, DENND2D, FCMR, LRRTM1, HPGDS, CTSS, TNFAIP8L2, ARHGAP25, S100A9, SGO2, MNDA, PYHIN1, CD200R1, PLB1, CCR1, GUCY1A1, TMEM144, CGAS, GPR85, BMPER, FABP5, TAGAP, SYK, ALDH1A1, CYBB, FOLR2, ASPG, C2, PLD4, ANPEP, C18orf54, MS4A7, MS4A14, NOD2, SNX20, DPEP2, ZNF528, LAIR1, ZNF558, TMC8, GSDMA, CX3CR1, RHOH, GFRA2, ROR2, P2RY12, PTAFR, RNASE6, GPR183, REPS2, ITGAM, CD14, SGCD, OSCAR, FPR1, GIMAP8, PIK3CD, P2RY6, S100Z, GPR82, GPR34, RGS19, GATM, RASGRP4, C3AR1, MBOAT1, CLEC7A, IL16, CTSW, KLHL6, SLFN11, CYSLTR1, PARP15, C1QB, C1QA, GPR160, NRROS, CTSF, TLR1, TLR6, SLCO2A1, ADGRE1, DES, LPL, B3GNT5, C20orf197, CD163, CD163L1, GPR150, ZNF366, ZC3H12D, PRSS36, CD28, FUCA1, RAB39A, HLA-DQB1, CIITA, FCER1A, LRRC3B, GPBAR1, TMEM150B, FAR2P1, HCLS1, ARHGAP45, TSPYL5, CCL13, P2RY13, FES, CSF1R, ACSM5, MACC1, UTY, ADAP2, CLDN6, APOBR, FMNL1, TNFAIP2, IKZF1, EVI2B, ZNF829, ZNF267, CD300LF, ZNF501, ARHGAP30, GPR141, FPR3, TLR5, CARD9, LRRK2, ANKRD34B, HLA-DRB1, SIRPB2, RYR1, POTEF, ZNF502, TLR7, HLA-DQA1, GM2A, AC079612.1, LAMB3, TMEM26, SIGLEC15, SERPINA1, C5AR1, MAP3K5, SPN, MYO5A, MPEG1, FAM49A, PLCG2, FCGR1B, SIRPA, ZNF248, SULT1C2, SLC29A3, HLA-DRB5, TUSC1, RCSD1, GRIN3A, ALPK2, GK, INPP5F, SUCNR1, RYR3, TOX, SFMBT2, CR1, FCGR3A, MAFB, GGTA1P, TMEM273, HLA-DOA, HLA-DMA, TAP2, HLA-DRA, AIF1, LST1, MICB, LILRB3, SLFN12L, CCDC144NL, DENND1C, TMEM200C, GIMAP1, LTC4S, GPSM3, SH3D21, LGMNP1, MS4A4E, HLA-DPB1, AC009948.1, LINC01857, HSPA7, AF127936.1, ITGB2- AS1, ZNF717, HLA-DRB6, HLA-DPA1, AL078590.3, CCDC144NL-AS1, ZNF736, NFAM1, GNAS-AS1, AC110995.1, HLA-DQA2, TNFSF12, L1TD1, PLEKHO2, LINC00996, HLA-DMB, LILRA6, FCGR2C, SOCS2-AS1, PCED1B-AS1, LINC00504, ASNSP1, FMN1, LINC01088, CARMN, AC116563.1, LINC01094, PCDHGA8, PCDHGB4, SIGLEC12, FPGT, SENCR, AC090559.1, AP002954.1, TIFAB, CLEC5A, AL133371.2, ITGB3, MRC1, PECAM1, MMP12, AC079062.1, AC138207.4, TXNIP, AP005131.6, AL132655.1, ADGRE4P, ZIM2-AS1, ZNF528-AS1, AC097634.1, HTATSF1P2, MILR1, CYFIP1, AL034397.3, CCL4, TUBGCP5, CCL4L2, CCL3L1, PIK3R6, PKD1L3, CCL3, TRG-AS1, ADORA3

## Protocol 1: 165 Down-regulated DEGs in CS52-C9n6-M microglia

ITGA3, SCIN, AASS, SEMA3B, SLC7A14, CLDN11, BIRC3, FAS, USP2, PHLPP2, RAB27B, FOXN3, OPN3, GPC1, MSANTD3, LRP6, CYBRD1, SEMA3C, CD82, BDKRB1, GPR143, PLP2, MMP15, CTSH, BBC3, DBP, ISYNA1, VIPR2, PIP5K1B, LGALS3BP, DTX4, PPM1H, GCNT2, CCNG1, LRRC42, KCNC4, DNAJC6, PLA2G4A, SLC2A1, SLC16A7, OGFRL1, NKX2-3, DUSP4, SLPI, PARD6B, KCNK15, CDKN1A, DOK4, BMP4, LRRN4, GLIS2, ARMCX1, LIF, FLNC, JUND, EDA2R, MTSS1L, DDB2, RAB11FIP5, MDM2, ADAMTS7, TFAP2A, LRRC1, PCSK6, SLC14A1, TP53, EPHA2, ADAM15, WNT9A, UBXN4, ALDH1L1, USP53, MYO10, GABRB2, SHROOM2, SYTL5, PTGES, DIXDC1, IL18, FAM177A1, WWC2, PLEKHH2, RETREG1, CXADR, VAV2, FDXR, AK4, PDPN, KIAA1522, WDR63, CAPN2, NIPAL1, PRKCI, SLC6A20, SPRY1, FAM160A1, SLC45A2, CMBL, FOXQ1, TP53INP1, ZNF219, PRKCB, STX3, SCG5, NSG1, EFNA1, FASN, SEMA3E, ELOVL6, SERPINB9, PPM1D, PDGFD, PYGO1, PKIA, BCL2L1, ZMAT3, PDP2, DHCR7, DAB1, SUSD5, HEG1, PHLDA3, FZD4, SEZ6L2, YPEL2, RPRM, PARD6G, CPNE7, LYNX1, KCNIP1, CTNNA3, NPW, FAM19A3, RPS27L, NAT8L, PPARA, NCR3LG1, IER5L, LINC01164, KAZN, CD55, NOS1AP, MUC19, NYNRIN, HLA- H, COL6A6, HLA-A, SSXP10, LINC00113, MIR34AHG, ANKRD18B, TMEM253, LNCTAM34A, C1orf226, PEG10, UPK3B, SPRR2F, PTCHD4, PURPL, PCDHGB6, AC124798.1, AC068700.1, AL157394.1, ANXA8, AL109976.1

## Protocol 1: 53 Up-regulated DEGs in CS29-C9n1-M microglia

GNA15, MAP3K13, HAS1, MAP3K8, EHHADH, ACOT2, RDH10, ARHGAP22, PUS7L, EDNRB, MNS1, ZNF300, TRIM4, CRIM1, VSIG4, SLAMF8, S100B, S100A9, CXCL5, CCDC68, ZNF667-AS1, ZNF528, IL7R, CD14, SPINK6, UTY, ACOT1, ARMCX2, ZNF829, ZNF471, ZNF677, ZNF311, IRAK4, ZNF667, ZNF568, INPP5F, ARMCX6, PCDHA2, AC016747.1, CEBPZOS, ZNF880, SVIL-AS1, ZNF717, ZNF736, GNAS-AS1, HLA-DQA2, ASNSP1, FPGT, ZNF350, FNBP1P1, ZNF790-AS1, ZNF595, ZNF229

## Protocol 1: 28 Down-regulated DEGs in CS29-C9n1-M microglia

CYP26B1, SOX8, GPM6B, SLC4A4, CDH7, SLC1A6, MYH14, ITGB6, PLPPR4, PLAGL1, EPCAM, ARHGAP40, KCNC3, LEFTY2, ADGRL3, FBXL21, PTCHD1, TMEM100, KIAA1024, SLN, HAS2, IGDCC3, TCIM, TMEM30B, PCDH9, SLITRK6, AC110619.1, PCDHB17P

## Protocol 2: 890 Up-regulated DEGs in CS52-C9n6-M microglia

SEMA3F, CYP26B1, TFPI, HOXA11, ITGA3, TMEM98, TMEM132A, BAIAP2L1, PROM1, TEAD3, AASS, NFIX, TENM1, TSPAN9, EHD3, GPRC5A, ISL1, CNTN1, WWTR1, SNAI2, ADGRA2, FHL1, EHD2, LRRC7, VCL, VCAN, TNC, EPHA3, DSG2, ADAMTS6, LTBP1, RCN1, DKK3, LAMC3, PTGER3, BCAR1, NNAT, CCDC85A, DCBLD2, PKP2, LAMC2, PTPRU, PRDM6, NCKAP1, CDH3, LIMCH1, ERBB3, MYLK, MPPED2, SYT1, PDZD4, FGFR3, ABCC9, PFN2, FSTL3, PTPN21, LMCD1, LIMS2, P4HA2, FCGR2B, CRMP1, FERMT2, GLI2, NOTCH3, TEAD2, ZNF532, CELSR1, CACNG4, MCAM, RAP1GAP, RARB, PPP1R12B, PAK3, FGFR1, ITGA8, FAP, NEBL, ADCY2, HOXA9, ITM2A, CLUL1, RUNX1T1, TNS1, MOXD1, MID2, COL5A3, KCNN2, COL4A4, DLG3, SERTAD4, FAT1, CTTN, WNT11, EPDR1, NOX4, GNAO1, NID2, ASAP3, DOCK9, ARHGAP28, SIGLEC1, CPXM1, HEPH, PITPNM2, LAMB1, MAP3K20, SEL1L3, ZFHX4, CCDC80, MYH7, SEMA6A, TYRO3, PHGDH, COL9A3, TGFB2, SORBS1, BAMBI, IL11, DSP, ABLIM1, PALM, KDELR3, RBFOX2, SIX4, EFS, NFATC4, PLTP, FERMT1, MYL9, JAG1, TBL1X, MID1, SRPX, PLS3, PCYT1B, KLHL4, FGD1, ZDHHC15, DRP2, NALCN, KLF5, FLT1, MEDAG, OLFM4, ZNF423, SALL1, CALB1, SFRP1, JPH1, NCALD, SH2D4A, NEFM, TUSC3, SYDE1, NUMBL, CLIP3, APLP1, MYH14, MEIS3, RAB3D, JAK3, TFPI2, CAV2, CAV1, MET, HOXA3, HOXA5, HOXA13, GRB10, FKBP14, CHCHD2, PTPRZ1, SERPINE1, SFRP4, GLI3, AEBP1, SLC1A1, PRUNE2, ASPN, ELAVL2, GLIS3, PDLIM1, GATA3, SPOCK2, DKK1, HOXB6, RAB34, CPE, AADAT, SH3D19, GLRB, B3GAT1, CCKBR, ARHGEF17, KIAA1549L, MDK, EXPH5, PPFIBP1, SYT10, LIN7A, KRT18, WNT5B, OAS3, OAS2, MGP, COL12A1, TPD52L1, CAP2, PERP, ARFGEF3, ADGRG6, PDE10A, MDFI, SMOC2, LAMA4, SEMA5A, C7, SPARC, RASGRF2, CDH6, NPR3, SLC27A6, LIFR, PDGFRB, STC2, AMOTL2, RBP1, PLSCR4, ITGB6, EVA1A, FN1, IGFBP2, IGFBP5, EFHD1, FHL2, QPCT, SDC1, EPHA4, MARK1, GPX7, ERRFI1, AKT3, ST6GALNAC5, ADGRL2, MFAP2, GBP1, ECE1, SLC2A1, CNN3, FILIP1, CTGF, PPL, KLF12, PCDH17, CCND2, C1orf198, LTBP2, KLHL29, PPP1R3C, HOXB3, TEK, SMAD9, GLT8D2, TBX2, ECHDC2, PLBD1, CAT, ADGRB2, CSMD2, INHBA, TWIST1, GLIPR2, KIAA1549, CALD1, CHST3, BICC1, NECAB1, NRK, PTGIS, HIF3A, ZNF391, SOX4, DOK4, EFNB2, BMP4, SOX9, BMP2, FLRT3, DSTN, LRRN4, MMP24, ID1, WNK4, GLIS2, RHOJ, PLEKHG3, AIF1L, TNFRSF19, GNG11, FGL2, SGCE, MGAT3, CDC42EP1, KRT17, PODXL, FOXP2, FLNC, LRRC17, CHN1, CGNL1, ISLR, LOXL1, PALLD, AJUBA, EGLN3, FGF13, PLPPR3, SH3BP4, DOCK6, LRCH2, PXDN, COL5A1, LAMA5, AKAP12, LILRB2, LRRC4B, GFPT2, MAP1B, RAI2, MATN2, SNAP25, NES, SYT11, PATJ, CHRM3, MYH10, DCLK1, EPSTI1, TRPC4, POSTN, MYH11, PDZD2, ZDHHC8P1, TEX15, LOXL2, TSPAN2, IAH1, FST, SOX5, DSC2, DSC3, FHOD3, COL4A2, DZIP1, ETS1, ANXA1, CEMIP2, PSAT1, CCNJL, MRAP2, EPHA7, KRT7, KCNK1, SERPINE2, CKAP4, FLNB, ADAMTS7, ABHD17C, ALPK3, SCN2A, GALNT5, DAB2IP, ANGPTL2, NOV, IL33, SULF1, YAP1, ITGA11, LRRC49, UACA, TTLL7, IFI44L, ARHGAP29, IFI44, EMILIN1, LOXL4, PLCE1, ADAMTS14, TET1, HECW2, CHRNA1, ITGAV, SLC40A1, PCDH10, PDE5A, FRAS1, SHROOM3, ANXA3, LEF1, EGF, FBN2, PIANP, AMIGO2, COL2A1, LGR5, VPS37B, NOVA1, FRMD6, FBLN5, DNAJA4, TPM1, ST8SIA2, TGFB1I1, ADAMTS18, CDH11, MYOCD, ADCYAP1, GATA6, GREB1L, P3H4, ERBB2, GRB7, FKBP10, IFITM3, FBN3, IGLON5, HSPG2, CYR61, TINAGL1, PTPRF, GPR161, RGS5, CRABP2, HMCN1, SYT14, VASH2, ATP8B2, LEFTY2, MBOAT2, ETNK2, ST6GAL2, AFF3, GALNT13, GULP1, CNTN4, RBMS3, OSBPL10, ITGA9, IL17RD, PHLDB2, BOC, ADAMTS16, OSMR, PAM, TENM2, TNFRSF21, DAAM2, SCUBE3, TPBG, EGFR, IGFBP3, NLGN4X, SHROOM2, DENND2A, TMEM47, SYTL5, SLC16A2, MAL2, VLDLR, NFIB, PLIN2, CDKN2B, NTRK2, PROSER2, FAM171A1, INA, GAS2, PAMR1, SERPING1, SERPINH1, JPH2, KIAA1755, MPP7, ITGB1, ADGRL3, DIXDC1, CRIM1, ANK3, ADAMTS12, CCDC3, ANO4, ASAP2, GFRA1, TMEM132D, TMEM56, PTPN14, TMEM178A, GRID2, SETBP1, TCF7L1, SPOCK1, PRDM8, RAB3C, ADGRA3, BMP6, PLA2R1, PTPRD, LURAP1L, PID1, KCTD15, DDAH1, CACNA2D1, SEMA3D, CHST9, ANGPT1, C4orf19, ENAH, SRSF12, NCAM2, ADAMTS1, ADAMTS5, TTN, FZD7, PXYLP1, GNA14, MMP16, KCNMA1, ADAMTS3, RWDD2B, PCDH1, GDF6, PPP2R2B, RBPMS, ERG, TSPAN18, CREB3L1, SLC37A3, DPYSL5, CLSTN2, RNF207, SHROOM4, SV2A, SIM2, TCHH, FNDC5, KALRN, CILP2, ABCG1, GPSM1, LY6E, COL26A1, DMKN, ITIH3, LRP5, AK4, SYNC, KIAA1522, MXRA8, MEGF6, DIRAS3, NFIA, NEXN, B3GALT2, NTNG1, GBP4, KCNT2, VANGL2, OLFML2B, CAPN2, ACTG2, ANTXR2, FSTL1, LMOD1, IGFBP7, FBLN2, PRICKLE2, APBB2, SGMS2, EMCN, PLXNB1, PRSS12, SCRG1, GUCY1A1, HHIP, ITGA2, EDIL3, F2RL2, GPX8, SEPT8, CGAS, FNDC1, DLC1, SBSPON, MAMDC2, TMEM246, PTCHD1, PCDH19, WNK2, SLITRK5, FAT3, SLC16A9, PKNOX2, DACT1, BEND7, FAM69B, JCAD, SALL2, TC2N, AMOTL1, GPR176, JAM3, ADAMTS15, FBN1, LARP6, GABRB3, DCHS1, CYB5R2, TUB, HDGFL3, PLEKHA7, NNMT, KIF7, RBPMS2, NAV2, MAP1A, CERCAM, IGF2, MYO5B, IGFBP6, ZNF558, LTBP3, SCARA3, VASN, KIF5C, BMP1, CAVIN2, SNTB2, AFAP1L2, CPT1C, PCSK9, MN1, EFNA1, CLIC4, CCDC8, LINGO1, PCDH7, CTNND2, FRMPD4, ZFPM2, IFFO2, RNF150, KRT8, CDH2, SGCD, GPR37, CAVIN3, PKIA, KRT19, DSEL, ENC1, FRMD5, TLN2, SYNPO, LAMB2, ID4, NEGR1, B3GALT1, CSDC2, RCAN2, ARNT2, SYNPO2, NIPAL4, PDE3A, EFEMP2, GXYLT2, RHOD, ADAMTS20, STOX2, NDNF, DAG1, PTPRM, CSPG4, HEG1, JUP, CBX2, SPTBN2, CTSF, MSRB3, CNTNAP2, SLCO2A1, FZD4, CD248, CSRP2, DPP10, LRRN1, TUBB6, LRRTM4, DMRTA1, RNF152, FOXC2, BDNF, SOX11, GCNT4, FIBIN, DPP7, FZD8, SAMD9L, PAWR, CAVIN1, SAMD12, CD163, AC098864.1, ADAMTSL1, KDELC2, CTXN1, RFLNA, AKAP5, SSC5D, TDRP, TSPYL5, ARSJ, F2R, PENK, MAB21L2, UNC5C, EXT1, FIGN, B4GALNT4, TSHZ2, BGN, PLCB1, ANXA2, PAPPA, CRIP2, COL18A1, TCEAL7, GJC1, GAS6, GPC6, GRIN2A, FBXL7, OPCML, LHFPL6, OLFML1, KIRREL1, ARSI, ST6GALNAC3, GPR173, PCDH9, PRKD1, EFNA5, SLITRK6, CLDN6, FAM110C, ROBO2, ROR1, PRKG1, NR2F2, LSAMP, AHNAK2, OLFML2A, C11orf87, KCNIP4, ZNF829, THBS2, CYP27C1, TEAD1, LYPD6, DCC, MAGI2, COL4A1, THSD4, PEAR1, C6orf132, COL4A5, AGRN, SBK1, COL25A1, S100A16, BEND4, KAZN, SPOCK3, FAT4, NUDT11, EPHB4, SLC6A9, AFAP1, MME, CACNA1H, ZNF502, FLNA, KIAA1671, KANK2, SVIL, STMN3, COL4A6, PARVA, FAM114A1, S100A10, HOXC6, TEAD4, ZNF677, ELOVL2, ZNF248, PEG3, ZNF568, TPM2, SH3BGRL2, PLN, TUSC1, RASSF9, FAM169A, ZNF521, ALPK2, DMD, TGM2, SAMD5, FCGR3A, FAM229B, RBM20, LINC00632, FOXO6, NHSL2, LAYN, FAM155A, GABBR1, CASC10, NYNRIN, DOK6, VGLL3, FIRRE, ITGA1, ARHGEF28, ISPD, TENM3, CEBPZOS, ZNF844, AL390729.1, HSPA7, MIAT, ERVMER34-1, C14orf132, HOTAIR, LINC01614, ZBED9, LINC00665, CCDC144NL-AS1, ZNF736, LINC00607, NR2F1-AS1, SHISA9, ARHGEF25, ANOS2P, PEG10, APELA, CARMN, SNHG18, AC125807.2, HOXA10, TRNP1, HOXA-AS2, MAGEL2, AP000662.1, FPGT, MEX3A, MIR100HG, TRIL, AL513534.1, AL353746.1, AL590004.3, TMEM178B, GATA6- AS1, BAHCC1, GABRQ, KCNQ1OT1, ZNF528-AS1, RASL10B, DOC2B, AC103702.2, ZBTB8B, CYFIP1, ARHGAP23, DACH1, NEFL, AC102945.2, AC145098.2, AC027290.2, MIR4697HG, AC092807.3, IQCJ-SCHIP1

## Protocol 2: 57 Down-regulated DEGs in CS52-C9n6-M microglia

ITGAL, STAB1, FAS, GNA15, KCNAB2, OSBPL3, IGF2BP2, IPCEF1, SLC1A3, LAT2, LIMK1, CD83, WNT5A, TNFRSF10B, TMIGD3, LRRC39, RPS4Y1, EDA2R, LGALS12, DDB2, MDM2, GYPC, CH25H, TP53, C1orf162, ZNF385B, USP6NL, LYPD1, VSIG4, LRRC58, MELTF, HTRA1, RAB3IL1, CD14, ZMAT3, RTTN, CD28, FCER1A, C9orf139, FUT7, TNFSF15, MAFF, RPS27L, INSIG1, ANKRD19P, PAX8-AS1, C5AR1, OGDHL, LRRC37A11P, SSXP10, MIR34AHG, GAS6-AS1, GPRC5D-AS1, PURPL, AC025569.1, FCGBP, AC079949.2

## Protocol 3: 3 Up-regulated DEGs in CS29-C9n1-M microglia

MAP3K13, IRAK4, FPGT

## Protocol 3: 1 Down-regulated DEG in CS29-C9n1-M microglia

SFTA1P

## References

Andrade-Talavera Y, Chen G, Pansieri J, Arroyo-García LE, Toleikis Z, Smirnovas V, Johansson J, Morozova-Roche L, Fisahn A. S100A9 amyloid growth and S100A9 fibril-induced impairment of gamma oscillations in area CA3 of mouse hippocampus ex vivo is prevented by Bri2 BRICHOS. Prog Neurobiol. 2022 Dec;219:102366. doi: 10.1016/j.pneurobio.2022.102366.

Appel SH, Beers DR, Zhao W. Amyotrophic lateral sclerosis is a systemic disease: peripheral contributions to inflammation-mediated neurodegeneration. Curr Opin Neurol. 2021 Oct 1;34(5):765–772. doi: 10.1097/WCO.0000000000000983.

Arterbery AS, Yao J, Ling A, Avitzur Y, Martinez M, Lobritto S, Deng Y, Geliang G, Mehta S, Wang G, Knight J, Ekong UD. Inflammasome Priming Mediated *via* Toll-Like Receptors 2 and 4, Induces Th1-Like Regulatory T Cells in *De Novo* Autoimmune Hepatitis. Front Immunol. 2018 Jul 19;9:1612. doi: 10.3389/fimmu.2018.01612.

Atanasio A, Decman V, White D, Ramos M, Ikiz B, Lee HC, Siao CJ, Brydges S, LaRosa E, Bai Y, Fury W, Burfeind P, Zamfirova R, Warshaw G, Orengo J, Oyejide A, Fralish M, Auerbach W, Poueymirou W, Freudenberg J, Gong G, Zambrowicz B, Valenzuela D, Yancopoulos G, Murphy A, Thurston G, Lai KM. C9orf72 ablation causes immune dysregulation characterized by leukocyte expansion, autoantibody production, and glomerulonephropathy in mice. Sci Rep. 2016 Mar 16;6:23204. doi: 10.1038/srep23204.

Babelova A, Moreth K, Tsalastra-Greul W, Zeng-Brouwers J, Eickelberg O, Young MF, Bruckner P, Pfeilschifter J, Schaefer RM, Gröne HJ, Schaefer L. Biglycan, a danger signal that activates the NLRP3 inflammasome via toll-like and P2X receptors. J Biol Chem. 2009 Sep 4;284(36):24035–48. doi: 10.1074/jbc.M109.014266.

Bai T, Chen CC, Lau LF. Matricellular protein CCN1 activates a proinflammatory genetic program in murine macrophages. J Immunol. 2010 Mar 15;184(6):3223–32. doi: 10.4049/jimmunol.0902792. Epub 2010 Feb 17.

Balendra R, Isaacs AM. C9orf72-mediated ALS and FTD: multiple pathways to disease. Nat Rev Neurol. 2018 Sep;14(9):544–558. doi: 10.1038/s41582-018-0047-2.

Bancroft EA, De La Mora M, Pandey G, Zarate SM, Srinivasan R. Extracellular S100B inhibits A-type voltage-gated potassium currents and increases L-type voltage-gated calcium channel activity in dopaminergic neurons. Glia. 2022 Dec;70(12):2330–2347. doi: 10.1002/glia.24254.

Banerjee P, Mehta AR, Nirujogi RS, Cooper J, James OG, Nanda J, Longden J, Burr K, McDade K, Salzinger A, Paza E, Newton J, Story D, Pal S, Smith C, Alessi DR, Selvaraj BT, Priller J, Chandran S. Cell-autonomous immune dysfunction driven by disrupted autophagy in *C9orf72*-ALS iPSC-derived microglia contributes to neurodegeneration. Sci Adv. 2023 Apr 21;9(16):eabq0651. doi: 10.1126/sciadv.abq0651.

Barnett KC, Li S, Liang K, Ting JP. A 360° view of the inflammasome: Mechanisms of activation, cell death, and diseases. Cell. 2023 May 25;186(11):2288–2312. doi: 10.1016/j.cell.2023.04.025.

Benson SA, Ernst JD. TLR2-dependent inhibition of macrophage responses to IFN-gamma is mediated by distinct, gene-specific mechanisms. PLoS One. 2009 Jul 24;4(7):e6329. doi: 10.1371/journal.pone.0006329.

Beers DR, Appel SH. Immune dysregulation in amyotrophic lateral sclerosis: mechanisms and emerging therapies. Lancet Neurol. 2019 Feb;18(2):211–220. doi: 10.1016/S1474-4422(18)30394-6.

Beers DR, Thonhoff JR, Faridar A, Thome AD, Zhao W, Wen S, Appel SH. Tregs Attenuate Peripheral Oxidative Stress and Acute Phase Proteins in ALS. Ann Neurol. 2022 Aug;92(2):195–200. doi: 10.1002/ana.26375.

Benatar M, Wuu J, Huey ED, McMillan CT, Petersen RC, Postuma R, McHutchison C, Dratch L, Arias JJ, Crawley A, Houlden H, McDermott MP, Cai X, Thakur N, Boxer A, Rosen H, Boeve BF, Dacks P, Cosentino S, Abrahams S, Shneider N, Lingor P, Shefner J, Andersen PM, Al-Chalabi A, Turner MR; Attendees of the Second International Pre-Symptomatic ALS Workshop. The Miami Framework for ALS and related neurodegenerative disorders: an integrated view of phenotype and biology. Nat Rev Neurol. 2024 Jun;20(6):364–376. doi: 10.1038/s41582-024-00961-z. Epub 2024 May 20.

Bi J, Dai J, Koivisto L, Larjava M, Bi L, Häkkinen L, Larjava H. Inflammasome and cytokine expression profiling in experimental periodontitis in the integrin β6 null mouse. Cytokine. 2019 Feb;114:135–142. doi: 10.1016/j.cyto.2018.11.011.

Bianchi R, Giambanco I, Donato R. S100B/RAGE-dependent activation of microglia via NF-kappaB and AP-1 Co-regulation of COX-2 expression by S100B, IL-1beta and TNF-alpha. Neurobiol Aging. 2010 Apr;31(4):665–77. doi: 10.1016/j.neurobiolaging.2008.05.017.

Bolger AM, Lohse M, Usadel B. Trimmomatic: a flexible trimmer for Illumina sequence data. Bioinformatics. 2014 Aug 1;30(15):2114–20. doi: 10.1093/bioinformatics/btu170.

Burberry A, Suzuki N, Wang JY, Moccia R, Mordes DA, Stewart MH, Suzuki-Uematsu S, Ghosh S, Singh A, Merkle FT, Koszka K, Li QZ, Zon L, Rossi DJ, Trowbridge JJ, Notarangelo LD, Eggan K. Loss-of-function mutations in the C9ORF72 mouse ortholog cause fatal autoimmune disease. Sci Transl Med. 2016 Jul 13;8(347):347ra93. doi: 10.1126/scitranslmed.aaf6038.

Butovsky O, Jedrychowski MP, Moore CS, Cialic R, Lanser AJ, Gabriely G, Koeglsperger T, Dake B, Wu PM, Doykan CE, Fanek Z, Liu L, Chen Z, Rothstein JD, Ransohoff RM, Gygi SP, Antel JP, Weiner HL. Identification of a unique TGF-β-dependent molecular and functional signature in microglia. Nat Neurosci. 2014 Jan;17(1):131–43. doi: 10.1038/nn.3599.

Cai X, Liang X, Wang K, Liu Y, Hao M, Li H, Dai X, Ding L. Pyroptosis-related lncRNAs: A novel prognosis signature of colorectal cancer. Front Oncol. 2022 Nov 30;12:983895. doi: 10.3389/fonc.2022.

Calma AD, Pavey N, Menon P, Vucic S. Neuroinflammation in amyotrophic lateral sclerosis: pathogenic insights and therapeutic implications. Curr Opin Neurol. 2024 Oct 1;37(5):585–592. doi: 10.1097/WCO.0000000000001279.

Cardenas AM, Sarlls JE, Kwan JY, Bageac D, Gala ZS, Danielian LE, Ray-Chaudhury A, Wang HW, Miller KL, Foxley S, Jbabdi S, Welsh RC, Floeter MK. Pathology of callosal damage in ALS: An *ex-vivo*, 7T diffusion tensor MRI study. Neuroimage Clin. 2017 Apr 30;15:200–208. doi: 10.1016/j.nicl.2017.04.024.

Chen, Y, Lun, ATL, and Smyth, GK. From reads to genes to pathways: differential expression analysis of RNA-Seq experiments using Rsubread and the edgeR quasi-likelihood pipeline. F1000Research 2016 5, 1438. doi: 10.12688/f1000research.8987.2.

Chen M, Geoffroy CG, Meves JM, Narang A, Li Y, Nguyen MT, Khai VS, Kong X, Steinke CL, Carolino KI, Elzière L, Goldberg MP, Jin Y, Zheng B. Leucine Zipper-Bearing Kinase Is a Critical Regulator of Astrocyte Reactivity in the Adult Mammalian CNS. Cell Rep. 2018 Mar 27;22(13):3587–3597. doi: 10.1016/j.celrep.2018.02.102.

Chen CX, Abdian N, Maussion G, Thomas RA, Demirova I, Cai E, Tabatabaei M, Beitel LK, Karamchandani J, Fon EA, Durcan TM. A Multistep Workflow to Evaluate Newly Generated iPSCs and Their Ability to Generate Different Cell Types. Methods Protoc. 2021 Jul 19;4(3):50. doi: 10.3390/mps4030050.

Chew J, Gendron TF, Prudencio M, Sasaguri H, Zhang YJ, Castanedes-Casey M, Lee CW, Jansen-West K, Kurti A, Murray ME, Bieniek KF, Bauer PO, Whitelaw EC, Rousseau L, Stankowski JN, Stetler C, Daughrity LM, Perkerson EA, Desaro P, Johnston A, Overstreet K, Edbauer D, Rademakers R, Boylan KB, Dickson DW, Fryer JD, Petrucelli L. Neurodegeneration. C9ORF72 repeat expansions in mice cause TDP-43 pathology, neuronal loss, and behavioral deficits. Science. 2015 Jun 5;348(6239):1151-4. doi: 10.1126/science.aaa9344.

Chi B, Öztürk MM, Paraggio CL, Leonard CE, Sanita ME, Dastpak M, O’Connell JD, Coady JA, Zhang J, Gygi SP, Lopez-Gonzalez R, Yin S, Reed R. Causal ALS genes impact the MHC class II antigen presentation pathway. Proc Natl Acad Sci U S A. 2023 Sep 26;120(39):e2305756120. doi: 10.1073/pnas.2305756120.

Costa DVS, Bon-Frauches AC, Silva AMHP, Lima-Júnior RCP, Martins CS, Leitão RFC, Freitas GB, Castelucci P, Bolick DT, Guerrant RL, Warren CA, Moura-Neto V, Brito GAC. 5- Fluorouracil Induces Enteric Neuron Death and Glial Activation During Intestinal Mucositis via a S100B-RAGE-NFκB-Dependent Pathway. Sci Rep. 2019 Jan 24;9(1):665. doi: 10.1038/s41598-018-36878-z.

Debye B, Schmülling L, Zhou L, Rune G, Beyer C, Johann S. Neurodegeneration and NLRP3 inflammasome expression in the anterior thalamus of SOD1(G93A) ALS mice. Brain Pathol. 2018 Jan;28(1):14–27. doi: 10.1111/bpa.12467.

de Calbiac H, Renault S, Haouy G, Jung V, Roger K, Zhou Q, Campanari ML, Chentout L, Demy DL, Marian A, Goudin N, Edbauer D, Guerrera C, Ciura S, Kabashi E. Poly-GP accumulation due to C9orf72 loss of function induces motor neuron apoptosis through autophagy and mitophagy defects. Autophagy. 2024 Oct;20(10):2164–2185. doi: 10.1080/15548627.2024.2358736.

DeJesus-Hernandez M, Mackenzie IR, Boeve BF, Boxer AL, Baker M, Rutherford NJ, Nicholson AM, Finch NA, Flynn H, Adamson J, Kouri N, Wojtas A, Sengdy P, Hsiung GY, Karydas A, Seeley WW, Josephs KA, Coppola G, Geschwind DH, Wszolek ZK, Feldman H, Knopman DS, Petersen RC, Miller BL, Dickson DW, Boylan KB, Graff-Radford NR, Rademakers R. Expanded GGGGCC hexanucleotide repeat in noncoding region of C9ORF72 causes chromosome 9p-linked FTD and ALS. Neuron. 2011 Oct 20;72(2):245–56. doi: 10.1016/j.neuron.2011.09.011.

Deora V, Lee JD, Albornoz EA, McAlary L, Jagaraj CJ, Robertson AAB, Atkin JD, Cooper MA, Schroder K, Yerbury JJ, Gordon R, Woodruff TM. The microglial NLRP3 inflammasome is activated by amyotrophic lateral sclerosis proteins. Glia. 2020 Feb;68(2):407–421. doi: 10.1002/glia.23728.

Dobin A, Davis CA, Schlesinger F, Drenkow J, Zaleski C, Jha S, Batut P, Chaisson M, Gingeras TR. STAR: ultrafast universal RNA-seq aligner. Bioinformatics. 2013 Jan 1;29(1):15–21. doi: 10.1093/bioinformatics/bts635.

Donnelly CJ, Zhang PW, Pham JT, Haeusler AR, Mistry NA, Vidensky S, Daley EL, Poth EM, Hoover B, Fines DM, Maragakis N, Tienari PJ, Petrucelli L, Traynor BJ, Wang J, Rigo F, Bennett CF, Blackshaw S, Sattler R, Rothstein JD. RNA toxicity from the ALS/FTD C9ORF72 expansion is mitigated by antisense intervention. Neuron. 2013 Oct 16;80(2):415–28. doi: 10.1016/j.neuron.2013.10.015.

Douvaras P, Sun B, Wang M, Kruglikov I, Lallos G, Zimmer M, Terrenoire C, Zhang B, Gandy S, Schadt E, Freytes DO, Noggle S, Fossati V. Directed Differentiation of Human Pluripotent Stem Cells to Microglia. Stem Cell Reports. 2017 Jun 6;8(6):1516–1524. doi: 10.1016/j.stemcr.2017.04.023.

Du X, Xu Y, Chen S, Fang M. Inhibited CSF1R Alleviates Ischemia Injury via Inhibition of Microglia M1 Polarization and NLRP3 Pathway. Neural Plast. 2020 Aug 28;2020:8825954. doi: 10.1155/2020/8825954.

Duggan MR, Peng Z, Sipilä PN, Lindbohm JV, Chen J, Lu Y, Davatzikos C, Erus G, Hohman TJ, Andrews SJ, Candia J, Tanaka T, Joynes CM, Alvarado CX, Nalls MA, Cordon J, Daya GN, An Y, Lewis A, Moghekar A, Palta P, Coresh J, Ferrucci L, Kivimäki M, Walker KA. Proteomics identifies potential immunological drivers of postinfection brain atrophy and cognitive decline. Nat Aging. 2024 Sep;4(9):1263–1278. doi: 10.1038/s43587-024-00682-4. Epub 2024 Aug 14.

Franz S, Ertel A, Engel KM, Simon JC, Saalbach A. Overexpression of S100A9 in obesity impairs macrophage differentiation via TLR4-NFkB-signaling worsening inflammation and wound healing. Theranostics. 2022 Jan 16;12(4):1659–1682. doi: 10.7150/thno.67174.

Freeman L, Guo H, David CN, Brickey WJ, Jha S, Ting JP. NLR members NLRC4 and NLRP3 mediate sterile inflammasome activation in microglia and astrocytes. J Exp Med. 2017 May 1;214(5):1351–1370. doi: 10.1084/jem.20150237.

Fu RH, Chen HJ, Hong SY. Glycine-Alanine Dipeptide Repeat Protein from C9-ALS Interacts with Sulfide Quinone Oxidoreductase (SQOR) to Induce the Activity of the NLRP3 Inflammasome in HMC3 Microglia: Irisflorentin Reverses This Interaction. Antioxidants (Basel). 2023a Oct 23;12(10):1896. doi: 10.3390/antiox12101896.

Fu J, Wu H. Structural Mechanisms of NLRP3 Inflammasome Assembly and Activation. Annu Rev Immunol. 2023b Apr 26;41:301–316. doi: 10.1146/annurev-immunol-081022-021207.

Gao C, Shi Q, Pan X, Chen J, Zhang Y, Lang J, Wen S, Liu X, Cheng TL, Lei K. Neuromuscular organoids model spinal neuromuscular pathologies in C9orf72 amyotrophic lateral sclerosis. Cell Rep. 2024 Mar 26;43(3):113892. doi: 10.1016/j.celrep.2024.113892.

Gendron TF, Petrucelli L. Disease Mechanisms of *C9ORF72* Repeat Expansions. Cold Spring Harb Perspect Med. 2018 Apr 2;8(4):a024224. doi: 10.1101/cshperspect.a024224.

Goutman SA, Hardiman O, Al-Chalabi A, Chió A, Savelieff MG, Kiernan MC, Feldman EL. Emerging insights into the complex genetics and pathophysiology of amyotrophic lateral sclerosis. Lancet Neurol. 2022 May;21(5):465–479. doi: 10.1016/S1474-4422(21)00414-2.

Grottelli S, Mezzasoma L, Scarpelli P, Cacciatore I, Cellini B, Bellezza I. Cyclo(His-Pro) inhibits NLRP3 inflammasome cascade in ALS microglial cells. Mol Cell Neurosci. 2019 Jan;94:23–31. doi: 10.1016/j.mcn.2018.11.002.

Gu L, Chen H, Geng R, Liang T, Chen Y, Wang Z, Ye L, Sun M, Shi Q, Wan G, Chang J, Wei J, Ma W, Xiao J, Bao X, Wang R. Endothelial pyroptosis-driven microglial activation in choroid plexus mediates neuronal apoptosis in hemorrhagic stroke rats. Neurobiol Dis. 2024 Oct 15;201:106695. doi: 10.1016/j.nbd.2024.106695.

Gugliandolo A, Giacoppo S, Bramanti P, Mazzon E. NLRP3 Inflammasome Activation in a Transgenic Amyotrophic Lateral Sclerosis Model. Inflammation. 2018 Feb;41(1):93–103. doi: 10.1007/s10753-017-0667-5.

Haenseler W, Sansom SN, Buchrieser J, Newey SE, Moore CS, Nicholls FJ, Chintawar S, Schnell C, Antel JP, Allen ND, Cader MZ, Wade-Martins R, James WS, Cowley SA. A Highly Efficient Human Pluripotent Stem Cell Microglia Model Displays a Neuronal-Co-culture-Specific Expression Profile and Inflammatory Response. Stem Cell Reports. 2017 Jun 6;8(6):1727–1742. doi: 10.1016/j.stemcr.2017.05.017.

He A, Liu Y, Zhang R, Mao Y, Liu W. CircSFMBT2-OA alleviates chondrocyte apoptosis and extracellular matrix degradation through repressing NF-κB/NLRP3 inflammasome activation. Heliyon. 2023 Jun 16;9(6):e17312. doi: 10.1016/j.heliyon.2023.e17312.

Hermani A, De Servi B, Medunjanin S, Tessier PA, Mayer D. S100A8 and S100A9 activate MAP kinase and NF-kappaB signaling pathways and trigger translocation of RAGE in human prostate cancer cells. Exp Cell Res. 2006 Jan 15;312(2):184–97. doi: 10.1016/j.yexcr.2005.10.013.

Herzog S, Fragkou PC, Arneth BM, Mkhlof S, Skevaki C. Myeloid CD169/Siglec1: An immunoregulatory biomarker in viral disease. Front Med (Lausanne). 2022 Sep 23;9:979373. doi: 10.3389/fmed.2022.979373.

Hickman SE, Kingery ND, Ohsumi TK, Borowsky ML, Wang LC, Means TK, El Khoury J. The microglial sensome revealed by direct RNA sequencing. Nat Neurosci. 2013 Dec;16(12):1896–905. doi: 10.1038/nn.3554.

Ho R, Workman MJ, Mathkar P, Wu K, Kim KJ, O’Rourke JG, Kellogg M, Montel V, Banuelos MG, Arogundade OA, Diaz-Garcia S, Oheb D, Huang S, Khrebtukova I, Watson L, Ravits J, Taylor K, Baloh RH, Svendsen CN. Cross-Comparison of Human iPSC Motor Neuron Models of Familial and Sporadic ALS Reveals Early and Convergent Transcriptomic Disease Signatures. Cell Syst. 2021 Feb 17;12(2):159–175.e9. doi: 10.1016/j.cels.2020.10.010.

Hong G, Zhang W, Li H, Shen X, Guo Z. Separate enrichment analysis of pathways for up- and downregulated genes. J R Soc Interface. 2013 Dec 18;11(92):20130950. doi: 10.1098/rsif.2013.0950.

Hou A, Lan W, Law KP, Khoo SC, Tin MQ, Lim YP, Tong L. Evaluation of global differential gene and protein expression in primary Pterygium: S100A8 and S100A9 as possible drivers of a signaling network. PLoS One. 2014 May 13;9(5):e97402. doi: 10.1371/journal.pone.0097402.

Huang da W, Sherman BT, Lempicki RA. Systematic and integrative analysis of large gene lists using DAVID bioinformatics resources. Nat Protoc. 2009(a);4(1):44–57. doi: 10.1038/nprot.2008.211.

Huang da W, Sherman BT, Lempicki RA. Bioinformatics enrichment tools: paths toward the comprehensive functional analysis of large gene lists. Nucleic Acids Res. 2009(b) Jan;37(1):1–13. doi: 10.1093/nar/gkn923.

Huang X, Feng Z, Jiang Y, Li J, Xiang Q, Guo S, Yang C, Fei L, Guo G, Zheng L, Wu Y, Chen Y. VSIG4 mediates transcriptional inhibition of *Nlrp3* and *Il-1*β in macrophages. Sci Adv. 2019 Jan 9;5(1):eaau7426. doi: 10.1126/sciadv.aau7426.

Ip WK, Medzhitov R. Macrophages monitor tissue osmolarity and induce inflammatory response through NLRP3 and NLRC4 inflammasome activation. Nat Commun. 2015 May 11;6:6931. doi: 10.1038/ncomms7931.

Izrael M, Slutsky SG, Revel M. Rising Stars: Astrocytes as a Therapeutic Target for ALS Disease. Front Neurosci. 2020 Jul 28;14:824. doi: 10.3389/fnins.2020.00824.

Ji H, Liu Z, Wang N, Jin J, Zhang J, Dong J, Wang F, Yan X, Gong Q, Zhao H, Sun H, Li Y, Hu S, You C. Integrated genomic, transcriptomic, and epigenetic analyses identify a leukotriene synthesis-related M2 macrophage gene signature that predicts prognosis and treatment vulnerability in gliomas. Front Immunol. 2022 Sep 8;13:970702. doi: 10.3389/fimmu.2022.970702.

Jin D, Dai Z, Zhao L, Ma T, Ma Y, Zhang Z. CYR61 is Involved in Neonatal Hypoxic-ischemic Brain Damage Via Modulating Astrocyte-mediated Neuroinflammation. Neuroscience. 2024 Aug 6;552:54–64. doi: 10.1016/j.neuroscience.2024.06.001.

Jun JI, Lau LF. CCN1 is an opsonin for bacterial clearance and a direct activator of Toll-like receptor signaling. Nat Commun. 2020 Mar 6;11(1):1242. doi: 10.1038/s41467-020-15075-5.

Kim JK, Choi EM, Shin HI, Kim CH, Hwang SH, Kim SM, Kwon BS. Characterization of monoclonal antibody specific to the Z39Ig protein, a member of immunoglobulin superfamily. Immunol Lett. 2005 Jul 15;99(2):153–61. doi: 10.1016/j.imlet.2005.02.012.

Kim TW, Staschke K, Bulek K, Yao J, Peters K, Oh KH, Vandenburg Y, Xiao H, Qian W, Hamilton T, Min B, Sen G, Gilmour R, Li X. A critical role for IRAK4 kinase activity in Toll-like receptor-mediated innate immunity. J Exp Med. 2007 May 14;204(5):1025–36. doi: 10.1084/jem.20061825.

Kim YC, Lee SE, Kim SK, Jang HD, Hwang I, Jin S, Hong EB, Jang KS, Kim HS. Toll-like receptor mediated inflammation requires FASN-dependent MYD88 palmitoylation. Nat Chem Biol. 2019 Sep;15(9):907–916. doi: 10.1038/s41589-019-0344-0.

Kirola L, Mukherjee A, Mutsuddi M. Recent Updates on the Genetics of Amyotrophic Lateral Sclerosis and Frontotemporal Dementia. Mol Neurobiol. 2022 Sep;59(9):5673–5694. doi: 10.1007/s12035-022-02934-z.

Lajqi T, Lang GP, Haas F, Williams DL, Hudalla H, Bauer M, Groth M, Wetzker R, Bauer R. Memory-Like Inflammatory Responses of Microglia to Rising Doses of LPS: Key Role of PI3Kγ. Front Immunol. 2019 Nov 8;10:2492. doi: 10.3389/fimmu.2019.02492.

Lall D, Lorenzini I, Mota TA, Bell S, Mahan TE, Ulrich JD, Davtyan H, Rexach JE, Muhammad AKMG, Shelest O, Landeros J, Vazquez M, Kim J, Ghaffari L, O’Rourke JG, Geschwind DH, Blurton-Jones M, Holtzman DM, Sattler R, Baloh RH. C9orf72 deficiency promotes microglial-mediated synaptic loss in aging and amyloid accumulation. Neuron. 2021 Jul 21;109(14):2275–2291.e8. doi: 10.1016/j.neuron.2021.05.020.

Lant SB, Robinson AC, Thompson JC, Rollinson S, Pickering-Brown S, Snowden JS, Davidson YS, Gerhard A, Mann DM. Patterns of microglial cell activation in frontotemporal lobar degeneration. Neuropathol Appl Neurobiol. 2014 Oct;40(6):686–96. doi: 10.1111/nan.12092.

Leaf IA, Nakagawa S, Johnson BG, Cha JJ, Mittelsteadt K, Guckian KM, Gomez IG, Altemeier WA, Duffield JS. Pericyte MyD88 and IRAK4 control inflammatory and fibrotic responses to tissue injury. J Clin Invest. 2017 Jan 3;127(1):321–334. doi: 10.1172/JCI87532.

Lee YL, Lin SK, Hong CY, Wang JS, Yang H, Lai EH, Chen MH, Kok SH. Major histocompatibility complex class II transactivator inhibits cysteine-rich 61 expression in osteoblastic cells and its implication in the pathogenesis of periapical lesions. J Endod. 2010 Jun;36(6):1021–5. doi: 10.1016/j.joen.2010.03.009.

Li B, Dewey CN. RSEM: accurate transcript quantification from RNA-Seq data with or without a reference genome. BMC Bioinformatics. 2011 Aug 4;12:323. doi: 10.1186/1471-2105-12-323.

Li S, Jiang L, Yang Y, Cao J, Zhang Q, Zhang J, Wang R, Deng X, Li Y. Siglec1 enhances inflammation through miR-1260-dependent degradation of IκBα in COPD. Exp Mol Pathol. 2020 Apr;113:104398. doi: 10.1016/j.yexmp.2020.104398.

Li D, Yu L, Zi J, Du X, Yan X, Chen H, Wang L, Zheng C, Wang G, Zhang J, Jiang Y. SLAMF8 Disrupts Epithelial Barrier in Chronic Rhinosinusitis with Nasal Polyps via M1 Macrophage Polarization. Ann Allergy Asthma Immunol. 2025 Jan 25:S1081–1206(25)00047-X. doi: 10.1016/j.anai.2025.01.020.

Lopez-Gonzalez R, Lu Y, Gendron TF, Karydas A, Tran H, Yang D, Petrucelli L, Miller BL, Almeida S, Gao FB. Poly(GR) in C9ORF72-Related ALS/FTD Compromises Mitochondrial Function and Increases Oxidative Stress and DNA Damage in iPSC-Derived Motor Neurons. Neuron. 2016 Oct 19;92(2):383–391. doi: 10.1016/j.neuron.2016.09.015.

Liu X, Wang J, Jin J, Hu Q, Zhao T, Wang J, Gao J, Man J. S100A9 deletion in microglia/macrophages ameliorates brain injury through the STAT6/PPARγ pathway in ischemic stroke. CNS Neurosci Ther. 2024 Aug;30(8):e14881. doi: 10.1111/cns.14881.

Lorenzini I, Alsop E, Levy J, Gittings LM, Lall D, Rabichow BE, Moore S, Pevey R, Bustos LM, Burciu C, Bhatia D, Singer M, Saul J, McQuade A, Tzioras M, Mota TA, Logemann A, Rose J, Almeida S, Gao FB, Marks M, Donnelly CJ, Hutchins E, Hung ST, Ichida J, Bowser R, Spires-Jones T, Blurton-Jones M, Gendron TF, Baloh RH, Van Keuren-Jensen K, Sattler R. Moderate intrinsic phenotypic alterations in *C9orf72*ALS/FTD iPSC-microglia despite the presence of C9orf72 pathological features. Front Cell Neurosci. 2023 Jun 6;17:1179796. doi: 10.3389/fncel.2023.1179796.

Lu C, Liu J, Escames G, Yang Y, Wu X, Liu Q, Chen J, Song Y, Wang Z, Deng C, Acuña-Castroviejo D, Wang X. PIK3CG Regulates NLRP3/GSDMD-Mediated Pyroptosis in Septic Myocardial Injury. Inflammation. 2023 Dec;46(6):2416–2432. doi: 10.1007/s10753-023-01889-0.

Majewski S, Klein P, Boillée S, Clarke BE, Patani R. Towards an integrated approach for understanding glia in Amyotrophic Lateral Sclerosis. Glia. 2025 Mar;73(3):591–607. doi: 10.1002/glia.24622.

Martínez-Muriana A, Mancuso R, Francos-Quijorna I, Olmos-Alonso A, Osta R, Perry VH, Navarro X, Gomez-Nicola D, López-Vales R. CSF1R blockade slows the progression of amyotrophic lateral sclerosis by reducing microgliosis and invasion of macrophages into peripheral nerves. Sci Rep. 2016 May 13;6:25663. doi: 10.1038/srep25663.

Masaki M, Ikeda A, Shiraki E, Oka S, Kawasaki T. Mixed lineage kinase LZK and antioxidant protein-1 activate NF-kappaB synergistically. Eur J Biochem. 2003 Jan;270(1):76–83. doi: 10.1046/j.1432-1033.2003.03363.x.

McQuade A, Coburn M, Tu CH, Hasselmann J, Davtyan H, Blurton-Jones M. Development and validation of a simplified method to generate human microglia from pluripotent stem cells. Mol Neurodegener. 2018 Dec 22;13(1):67. doi: 10.1186/s13024-018-0297-x.

Mehta AR, Gregory JM, Dando O, Carter RN, Burr K, Nanda J, Story D, McDade K, Smith C, Morton NM, Mahad DJ, Hardingham GE, Chandran S, Selvaraj BT. Mitochondrial bioenergetic deficits in C9orf72 amyotrophic lateral sclerosis motor neurons cause dysfunctional axonal homeostasis. Acta Neuropathol. 2021 Feb;141(2):257–279. doi: 10.1007/s00401-020-02252-5.

Mimic S, Aru B, Pehlivanoğlu C, Sleiman H, Andjus PR, Yanıkkaya Demirel G. Immunology of amyotrophic lateral sclerosis - role of the innate and adaptive immunity. Front Neurosci. 2023 Nov 30;17:1277399. doi: 10.3389/fnins.2023.1277399.

Mooring M, Yeung GA, Luukkonen P, Liu S, Akbar MW, Zhang GJ, Balogun O, Yu X, Mo R, Nejak-Bowen K, Poyurovsky MV, Booth CJ, Konnikova L, Shulman GI, Yimlamai D. Hepatocyte CYR61 polarizes profibrotic macrophages to orchestrate NASH fibrosis. Sci Transl Med. 2023 Sep 27;15(715):eade3157. doi: 10.1126/scitranslmed.ade3157.

Muffat J, Li Y, Yuan B, Mitalipova M, Omer A, Corcoran S, Bakiasi G, Tsai LH, Aubourg P, Ransohoff RM, Jaenisch R. Efficient derivation of microglia-like cells from human pluripotent stem cells. Nat Med. 2016 Nov;22(11):1358–1367. doi: 10.1038/nm.4189.

Nicholson AM, Zhou X, Perkerson RB, Parsons TM, Chew J, Brooks M, DeJesus-Hernandez M, Finch NA, Matchett BJ, Kurti A, Jansen-West KR, Perkerson E, Daughrity L, Castanedes-Casey M, Rousseau L, Phillips V, Hu F, Gendron TF, Murray ME, Dickson DW, Fryer JD, Petrucelli L, Rademakers R. Loss of Tmem106b is unable to ameliorate frontotemporal dementia-like phenotypes in an AAV mouse model of C9ORF72-repeat induced toxicity. Acta Neuropathol Commun. 2018 May 31;6(1):42. doi: 10.1186/s40478-018-0545-x.

Nijs M, Van Damme P. The genetics of amyotrophic lateral sclerosis. Curr Opin Neurol. 2024 Oct 1;37(5):560–569. doi: 10.1097/WCO.0000000000001294.

Nunomura S, Uta D, Kitajima I, Nanri Y, Matsuda K, Ejiri N, Kitajima M, Ikemitsu H, Koga M, Yamamoto S, Honda Y, Takedomi H, Andoh T, Conway SJ, Izuhara K. Periostin activates distinct modules of inflammation and itching downstream of the type 2 inflammation pathway. Cell Rep. 2023 Jan 31;42(1):111933. doi: 10.1016/j.celrep.2022.111933.

O’Rourke JG, Bogdanik L, Yáñez A, Lall D, Wolf AJ, Muhammad AK, Ho R, Carmona S, Vit JP, Zarrow J, Kim KJ, Bell S, Harms MB, Miller TM, Dangler CA, Underhill DM, Goodridge HS, Lutz CM, Baloh RH. C9orf72 is required for proper macrophage and microglial function in mice. Science. 2016 Mar 18;351(6279):1324-9. doi: 10.1126/science.aaf1064.

Pappalardo XG, Jansen G, Amaradio M, Costanza J, Umeton R, Guarino F, De Pinto V, Oliver SG, Messina A, Nicosia G. Inferring gene regulatory networks of ALS from blood transcriptome profiles. Heliyon. 2024 Nov 26;10(23):e40696. doi: 10.1016/j.heliyon.2024.e40696.

Peferoen LA, Vogel DY, Ummenthum K, Breur M, Heijnen PD, Gerritsen WH, Peferoen-Baert RM, van der Valk P, Dijkstra CD, Amor S. Activation status of human microglia is dependent on lesion formation stage and remyelination in multiple sclerosis. J Neuropathol Exp Neurol. 2015 Jan;74(1):48–63. doi: 10.1097/NEN.0000000000000149.

Pruenster M, Immler R, Roth J, Kuchler T, Bromberger T, Napoli M, Nussbaumer K, Rohwedder I, Wackerbarth LM, Piantoni C, Hennis K, Fink D, Kallabis S, Schroll T, Masgrau- Alsina S, Budke A, Liu W, Vestweber D, Wahl-Schott C, Roth J, Meissner F, Moser M, Vogl T, Hornung V, Broz P, Sperandio M. E-selectin-mediated rapid NLRP3 inflammasome activation regulates S100A8/S100A9 release from neutrophils via transient gasdermin D pore formation. Nat Immunol. 2023 Dec;24(12):2021–2031. doi: 10.1038/s41590-023-01656-1.

Puck A, Künig S, Modak M, May L, Fritz P, Battin C, Radakovics K, Steinberger P, Reipert BM, Crowe BA, Stöckl J. The soluble cytoplasmic tail of CD45 regulates T-cell activation via TLR4 signaling. Eur J Immunol. 2021 Dec;51(12):3176–3185. doi: 10.1002/eji.202149227.

Qian J, Luo F, Yang J, Liu J, Liu R, Wang L, Wang C, Deng Y, Lu Z, Wang Y, Lu M, Wang JY, Chu Y. TLR2 Promotes Glioma Immune Evasion by Downregulating MHC Class II Molecules in Microglia. Cancer Immunol Res. 2018 Oct;6(10):1220–1233. doi: 10.1158/2326-6066.CIR-18-0020.

Qin W, Rong X, Yu C, Jia P, Yang J, Zhou G. Knockout of SLAMF8 attenuates collagen-induced rheumatoid arthritis in mice through inhibiting TLR4/NF-κB signaling pathway. Int Immunopharmacol. 2022 Jun;107:108644. doi: 10.1016/j.intimp.2022.108644.

Radian AD, Khare S, Chu LH, Dorfleutner A, Stehlik C. ATP binding by NLRP7 is required for inflammasome activation in response to bacterial lipopeptides. Mol Immunol. 2015 Oct;67(2 Pt B):294–302. doi: 10.1016/j.molimm.2015.06.013.

Rai AN, Thornton JA, Stokes J, Sunesara I, Swiatlo E, Nanduri B. Polyamine transporter in Streptococcus pneumoniae is essential for evading early innate immune responses in pneumococcal pneumonia. Sci Rep. 2016 Jun 1;6:26964. doi: 10.1038/srep26964.

Renton AE, Majounie E, Waite A, Simón-Sánchez J, Rollinson S, Gibbs JR, Schymick JC, Laaksovirta H, van Swieten JC, Myllykangas L, Kalimo H, Paetau A, Abramzon Y, Remes AM, Kaganovich A, Scholz SW, Duckworth J, Ding J, Harmer DW, Hernandez DG, Johnson JO, Mok K, Ryten M, Trabzuni D, Guerreiro RJ, Orrell RW, Neal J, Murray A, Pearson J, Jansen IE, Sondervan D, Seelaar H, Blake D, Young K, Halliwell N, Callister JB, Toulson G, Richardson A, Gerhard A, Snowden J, Mann D, Neary D, Nalls MA, Peuralinna T, Jansson L, Isoviita VM, Kaivorinne AL, Hölttä-Vuori M, Ikonen E, Sulkava R, Benatar M, Wuu J, Chiò A, Restagno G, Borghero G, Sabatelli M; ITALSGEN Consortium, Heckerman D, Rogaeva E, Zinman L, Rothstein JD, Sendtner M, Drepper C, Eichler EE, Alkan C, Abdullaev Z, Pack SD, Dutra A, Pak E, Hardy J, Singleton A, Williams NM, Heutink P, Pickering-Brown S, Morris HR, Tienari PJ, Traynor BJ. A hexanucleotide repeat expansion in C9ORF72 is the cause of chromosome 9p21-linked ALS-FTD. Neuron. 2011 Oct 20;72(2):257–68. doi: 10.1016/j.neuron.2011.09.010.

Riva M, Källberg E, Björk P, Hancz D, Vogl T, Roth J, Ivars F, Leanderson T. Induction of nuclear factor-κB responses by the S100A9 protein is Toll-like receptor-4-dependent. Immunology. 2012 Oct;137(2):172–82. doi: 10.1111/j.1365-2567.2012.03619.x.

Rivers-Auty J, Hoyle C, Pointer A, Lawrence C, Pickering-Brown S, Brough D, Ryan S. *C9orf72*dipeptides activate the NLRP3 inflammasome. Brain Commun. 2024 Aug 20;6(5):fcae282. doi: 10.1093/braincomms/fcae282.

Roedig H, Nastase MV, Frey H, Moreth K, Zeng-Brouwers J, Poluzzi C, Hsieh LT, Brandts C, Fulda S, Wygrecka M, Schaefer L. Biglycan is a new high-affinity ligand for CD14 in macrophages. Matrix Biol. 2019 Apr;77:4–22. doi: 10.1016/j.matbio.2018.05.006.

Sanfilippo C, Longo A, Lazzara F, Cambria D, Distefano G, Palumbo M, Cantarella A, Malaguarnera L, Di Rosa M. CHI3L1 and CHI3L2 overexpression in motor cortex and spinal cord of sALS patients. Mol Cell Neurosci. 2017 Dec;85:162–169. doi: 10.1016/j.mcn.2017.10.001.

Schwanekamp JA, Lorts A, Vagnozzi RJ, Vanhoutte D, Molkentin JD. Deletion of Periostin Protects Against Atherosclerosis in Mice by Altering Inflammation and Extracellular Matrix Remodeling. Arterioscler Thromb Vasc Biol. 2016 Jan;36(1):60–8. doi: 10.1161/ATVBAHA.115.306397.

Sharma BR, Kanneganti TD. NLRP3 inflammasome in cancer and metabolic diseases. Nat Immunol. 2021 May;22(5):550–559. doi: 10.1038/s41590-021-00886-5.

Sheng M, Weng Y, Cao Y, Zhang C, Lin Y, Yu W. Caspase 6/NR4A1/SOX9 signaling axis regulates hepatic inflammation and pyroptosis in ischemia-stressed fatty liver. Cell Death Discov. 2023 Mar 28;9(1):106. doi: 10.1038/s41420-023-01396-z.

Sheng M, Liu W, Cao Y, Wang S, Lin Y, Yu W. Targeting S100A9-TLR2 axis controls macrophage NLRP3 inflammasome activation in fatty liver ischemia reperfusion injury. Shock. 2024 Oct 21. doi: 10.1097/SHK.0000000000002470.

Shu X, Wei C, Tu WY, Zhong K, Qi S, Wang A, Bai L, Zhang SX, Luo B, Xu ZZ, Zhang K, Shen C. Negative regulation of TREM2-mediated C9orf72 poly-GA clearance by the NLRP3 inflammasome. Cell Rep. 2023 Feb 28;42(2):112133. doi: 10.1016/j.celrep.2023.112133.

Souza de Lima D, Nunes VCL, Ogusku MM, Sadahiro A, Pontillo A, Alencar BC. Polymorphisms in SIGLEC1 contribute to susceptibility to pulmonary active tuberculosis possibly through the modulation of IL-1ß. Infect Genet Evol. 2017 Nov;55:313–317. doi: 10.1016/j.meegid.2017.09.031.

Sudria-Lopez E, Koppers M, de Wit M, van der Meer C, Westeneng HJ, Zundel CA, Youssef SA, Harkema L, de Bruin A, Veldink JH, van den Berg LH, Pasterkamp RJ. Full ablation of C9orf72 in mice causes immune system-related pathology and neoplastic events but no motor neuron defects. Acta Neuropathol. 2016 Jul;132(1):145–7. doi: 10.1007/s00401-016-1581-x.

Suzuki N, Suzuki S, Duncan GS, Millar DG, Wada T, Mirtsos C, Takada H, Wakeham A, Itie A, Li S, Penninger JM, Wesche H, Ohashi PS, Mak TW, Yeh WC. Severe impairment of interleukin-1 and Toll-like receptor signalling in mice lacking IRAK-4. Nature. 2002 Apr 18;416(6882):750–6. doi: 10.1038/nature736.

Tang YM, Pulimood NS, Stifani S. Comparing the Characteristics of Microglia Preparations Generated Using Different Human iPSC-Based Differentiation Methods to Model Neurodegenerative Diseases. ASN Neuro. 2022 Jan-Dec;14:17590914221145105. doi: 10.1177/17590914221145105.

Thiry L, Clément JP, Haag R, Kennedy TE, Stifani S. Optimization of Long-Term Human iPSC-Derived Spinal Motor Neuron Culture Using a Dendritic Polyglycerol Amine-Based Substrate. ASN Neuro. 2022 Jan-Dec;14:17590914211073381. doi: 10.1177/17590914211073381.

Thiry L, Hamel R, Pluchino S, Durcan T, Stifani S. Characterization of Human iPSC-derived Spinal Motor Neurons by Single-cell RNA Sequencing. Neuroscience. 2020 Dec 1;450:57–70. doi: 10.1016/j.neuroscience.2020.04.041.

Thiry L, Sirois J, Durcan TM, Stifani S. Generation of human iPSC-derived phrenic-like motor neurons to model respiratory motor neuron degeneration in ALS. Commun Biol. 2024 Feb 28;7(1):238. doi: 10.1038/s42003-024-05925-z.

Trageser KJ, Yang EJ, Smith C, Iban-Arias R, Oguchi T, Sebastian-Valverde M, Iqbal UH, Wu H, Estill M, Al Rahim M, Raval U, Herman FJ, Zhang YJ, Petrucelli L, Pasinetti GM. Inflammasome-Mediated Neuronal-Microglial Crosstalk: a Therapeutic Substrate for the Familial C9orf72 Variant of Frontotemporal Dementia/Amyotrophic Lateral Sclerosis. Mol Neurobiol. 2023 Jul;60(7):4004–4016. doi: 10.1007/s12035-023-03315-w.

Tran H, Almeida S, Moore J, Gendron TF, Chalasani U, Lu Y, Du X, Nickerson JA, Petrucelli L, Weng Z, Gao FB. Differential Toxicity of Nuclear RNA Foci versus Dipeptide Repeat Proteins in a Drosophila Model of C9ORF72 FTD/ALS. Neuron. 2015 Sep 23;87(6):1207–1214. doi: 10.1016/j.neuron.2015.09.015.

Unamuno X, Gómez-Ambrosi J, Ramírez B, Rodríguez A, Becerril S, Valentí V, Moncada R, Silva C, Salvador J, Frühbeck G, Catalán V. NLRP3 inflammasome blockade reduces adipose tissue inflammation and extracellular matrix remodeling. Cell Mol Immunol. 2021 Apr;18(4):1045–1057. doi: 10.1038/s41423-019-0296-z.

Vahsen BF, Nalluru S, Morgan GR, Farrimond L, Carroll E, Xu Y, Cramb KML, Amein B, Scaber J, Katsikoudi A, Candalija A, Carcolé M, Dafinca R, Isaacs AM, Wade-Martins R, Gray E, Turner MR, Cowley SA, Talbot K. C9orf72-ALS human iPSC microglia are pro-inflammatory and toxic to co-cultured motor neurons via MMP9. Nat Commun. 2023 Sep 22;14(1):5898. doi: 10.1038/s41467-023-41603-0.

Vasudevan SO, Russo AJ, Kumari P, Vanaja SK, Rathinam VA. A TLR4-independent critical role for CD14 in intracellular LPS sensing. Cell Rep. 2022 May 3;39(5):110755. doi: 10.1016/j.celrep.2022.110755.

Vidmar L, Maver A, Drulović J, Sepčić J, Novaković I, Ristič S, Šega S, Peterlin B. Multiple Sclerosis patients carry an increased burden of exceedingly rare genetic variants in the inflammasome regulatory genes. Sci Rep. 2019 Jun 24;9(1):9171. doi: 10.1038/s41598-019-45598-x.

Villarreal A, Aviles Reyes RX, Angelo MF, Reines AG, Ramos AJ. S100B alters neuronal survival and dendrite extension via RAGE-mediated NF-κB signaling. J Neurochem. 2011 Apr;117(2):321–32. doi: 10.1111/j.1471-4159.2011.07207.x.

Villarreal A, Seoane R, González Torres A, Rosciszewski G, Angelo MF, Rossi A, Barker PA, Ramos AJ. S100B protein activates a RAGE-dependent autocrine loop in astrocytes: implications for its role in the propagation of reactive gliosis. J Neurochem. 2014 Oct;131(2):190–205. doi: 10.1111/jnc.12790.

Vizuete AFK, Fróes F, Seady M, Zanotto C, Bobermin LD, Roginski AC, Wajner M, Quincozes-Santos A, Gonçalves CA. Early effects of LPS-induced neuroinflammation on the rat hippocampal glycolytic pathway. J Neuroinflammation. 2022 Oct 11;19(1):255. doi: 10.1186/s12974-022-02612-w.

Wang Y, Ding J, Song H, Teng Y, Fang X. VSIG4 regulates macrophages polarization and alleviates inflammation through activating PI3K/AKT and inhibiting TLR4/NF-κB pathway in myocardial ischemia-reperfusion injury rats. Physiol Int. 2022a Sep 5. doi: 10.1556/2060.2022.00055.

Wang Y, Jin Z, Sun J, Chen X, Xie P, Zhou Y, Wang S. The role of activated monocyte IFN/SIGLEC1 signalling in Graves’ disease. J Endocrinol. 2022b Sep 5;255(1):1–9. doi: 10.1530/JOE-21-0453.

Wang G, Huang K, Tian Q, Guo Y, Liu C, Li Z, Yu Z, Zhang Z, Li M. S100A9 aggravates early brain injury after subarachnoid hemorrhage via inducing neuroinflammation and inflammasome activation. iScience. 2024 Feb 9;27(3):109165. doi: 10.1016/j.isci.2024.109165.

Xie Y, Luo X, He H, Tang M. Novel Insight Into the Role of Immune Dysregulation in Amyotrophic Lateral Sclerosis Based on Bioinformatic Analysis. Front Neurosci. 2021 Apr 30;15:657465. doi: 10.3389/fnins.2021.657465.

Yan P, Zhu A, Liao F, Xiao Q, Kraft A, Gonzales E, Perez R, Greenberg SM, Holtzman D, Lee JM. Minocycline reduces spontaneous hemorrhage in mouse models of cerebral amyloid angiopathy. Stroke. 2015 Jun;46(6):1633–1640. doi: 10.1161/STROKEAHA.115.008582.

Yao L, Song J, Meng XW, Ge JY, Du BX, Yu J, Ji FH. Periostin aggravates NLRP3 inflammasome-mediated pyroptosis in myocardial ischemia-reperfusion injury. Mol Cell Probes. 2020 Oct;53:101596. doi: 10.1016/j.mcp.2020.101596.

Yi K, Cui X, Liu X, Wang Y, Zhao J, Yang S, Xu C, Yang E, Xiao M, Hong B, Fang C, Kang C, Tan Y, Wang Q. PTRF/Cavin-1 as a Novel RNA-Binding Protein Expedites the NF-κB/PD- L1 Axis by Stabilizing lncRNA NEAT1, Contributing to Tumorigenesis and Immune Evasion in Glioblastoma. Front Immunol. 2022 Jan 6;12:802795. doi: 10.3389/fimmu.2021.802795.

Zeng X, Liu G, Peng W, He J, Cai C, Xiong W, Chen S, Yang M, Dong Z. Combined deficiency of SLAMF8 and SLAMF9 prevents endotoxin-induced liver inflammation by downregulating TLR4 expression on macrophages. Cell Mol Immunol. 2020 Feb;17(2):153–162. doi: 10.1038/s41423-018-0191-z.

Zhang Y, Chen K, Sloan SA, Bennett ML, Scholze AR, O’Keeffe S, Phatnani HP, Guarnieri P, Caneda C, Ruderisch N, Deng S, Liddelow SA, Zhang C, Daneman R, Maniatis T, Barres BA, Wu JQ. An RNA-sequencing transcriptome and splicing database of glia, neurons, and vascular cells of the cerebral cortex. J Neurosci. 2014 Sep 3;34(36):11929–47. doi: 10.1523/JNEUROSCI.1860-14.2014.

Zhang D, Shen X, Pang K, Yang Z, Yu A. VSIG4 alleviates intracerebral hemorrhage induced brain injury by suppressing TLR4-regulated inflammatory response. Brain Res Bull. 2021 Nov;176:67–75. doi: 10.1016/j.brainresbull.2021.08.008.

Zhao W, Beers DR, Henkel JS, Zhang W, Urushitani M, Julien JP, Appel SH. Extracellular mutant SOD1 induces microglial-mediated motoneuron injury. Glia. 2010 Jan 15;58(2):231–43. doi: 10.1002/glia.20919.

Zhao W, Beers DR, Bell S, Wang J, Wen S, Baloh RH, Appel SH. TDP-43 activates microglia through NF-κB and NLRP3 inflammasome. Exp Neurol. 2015 Nov;273:24–35. doi: 10.1016/j.expneurol.2015.07.019.

Zhao T, Zhang X, Cui X, Su S, Li L, Chen Y, Wang N, Sun L, Zhao J, Zhang J, Han X, Cao J. Inhibiting the IRAK4/NF-κB/NLRP3 signaling pathway can reduce pyroptosis in hippocampal neurons and seizure episodes in epilepsy. Exp Neurol. 2024 Jul;377:114794. doi: 10.1016/j.expneurol.2024.114794.

Zheng Y, Lee S, Liang X, Wei S, Moon HG, Jin Y. Suppression of PTRF alleviates the polymicrobial sepsis induced by cecal ligation and puncture in mice. J Infect Dis. 2013 Dec 1;208(11):1803–12. doi: 10.1093/infdis/jit364.

Zheng M, Li Z, Feng Y, Zhang X. CD14 and CSF1R as developmental molecular targets for the induction of osteoarthritis. Int J Clin Exp Pathol. 2023 Aug 15;16(8):184–198.

Zhong J, Wang C, Zhang D, Yao X, Zhao Q, Huang X, Lin F, Xue C, Wang Y, He R, Li XY, Li Q, Wang M, Zhao S, Afridi SK, Zhou W, Wang Z, Xu Y, Xu Z. PCDHA9 as a candidate gene for amyotrophic lateral sclerosis. Nat Commun. 2024 Mar 11;15(1):2189. doi: 10.1038/s41467-024-46333-5.

Zhou HH, Zhang YM, Zhang SP, Xu QX, Tian YQ, Li P, Cao D, Zheng YQ. Suppression of PTRF Alleviates Post-Infectious Irritable Bowel Syndrome via Downregulation of the TLR4 Pathway in Rats. Front Pharmacol. 2021 Oct 7;12:724410. doi: 10.3389/fphar.2021.724410.

Zhu D, Chen S, Sheng P, Wang Z, Li Y, Kang X. POSTN promotes nucleus pulposus cell senescence and extracellular matrix metabolism via activing Wnt/β-catenin and NF-κB signal pathway in intervertebral disc degeneration. Cell Signal. 2024 Sep;121:111277. doi: 10.1016/j.cellsig.2024.111277.

